# Bone diagenesis in Iron Age Siberia: a histological and microtomographic study of Tunnug 1 (Russia, 2^nd^–4^th^ c. CE)

**DOI:** 10.64898/2026.09.09.749369

**Authors:** Lolita Trenchat, Gino Caspari, Marco Milella, Timur Sadykov, Jegor Blochin, Sam C. Lin, Nicolas Vanderesse, Eline M.J. Schotsmans

## Abstract

The Siberian site of Tunnug 1, located in the Uyuk Valley (Tuva Republic, Russia), encompasses an Early Scythian burial mound (9th century BCE) and a peripheral cemetery attributed to the Kokel culture (2nd–4th centuries CE). Situated within a permafrost-affected environment, the site offers an opportunity to investigate bone preservation under long-term freeze-thaw cycle conditions and explore the relationship between funerary treatment and bone diagenesis. This study presents a combined histological and microtomographic analysis of 76 bone samples from 71 individuals across 36 burials. Transmitted light microscopy, scanning electron microscopy (SEM) and micro-computed tomography (micro-CT) were employed to document biological, chemical and physical degradation. Bone preservation was evaluated through three approaches: the traditional Oxford Histological Index (OHI), quantitative image analysis and assessment of (micro)cracking. The results revealed moderate bone preservation (OHI 2-4). Biological degradation in the form of Microscopical Focal Destruction (MFD) and chemical alteration were identified in six individuals. Enlarged canaliculi were observed in every bone sample. Microcracks were present in all individuals, visible exclusively under SEM. No clear diagenetic signatures linked to specific funerary practices were identified, though variations in preservation suggest a possible influence of burial depth. The distinct impact of permafrost and long-term freeze–thaw cycles on bone microstructure could not be isolated, underscoring the multifactorial nature of diagenesis. This study highlights the complementary value of combining multiple analytical methods assessing bone diagenesis and emphasises the need for continued experimental research into the effects of freezing environments on human remains.

## 1. Introduction

Histotaphonomic analysis of bone microstructure is used in archaeology to investigate diagenesis in relation to depositional environments (Dal Sasso et al. 2014; Booth et al. 2015; Papakonstantinou et al. 2020) and, where the evidence allows, to reconstruct mortuary practices (e.g. Trenchat et al. 2025). Bone diagenesis refers to the physical, chemical and biological alterations affecting skeletal material after death (Bell et al. 1991; Nielsen-Marsh et al. 2007; Smith et al. 2007). In archaeological contexts, the most commonly documented form of bone degradation is bacterial alteration, known as Microscopical Focal Destruction (MFD). Hackett (1981) classified them, using transmitted light microscopy, into three morphological types: budded, lamellate and linear-longitudinal. However, no data are currently available on the possible distinct biological origins of these morphological differences (Turner-Walker 2023). Wedl (1864) provided the first description of tunnels of degradation, subsequently named Wedl tunnelling. Initially attributed to fungal activity (Marchiafava et al. 1974), these alterations are now recognised as the product of cyanobacterial colonisation (Turner-Walker 2019; Eriksen et al. 2020; Turner-Walker et al. 2023). The question of whether the bacteria responsible for these alterations are exogenous, deriving from the surrounding soil, or endogenous, originating from the gut microbiome of the deceased, has been the subject of considerable debate in the literature (e.g. White and Booth 2014; Booth and Madgwick 2016; Brönnimann et al. 2018; Turner-Walker et al. 2023). However, a growing consensus is emerging that both sources may contribute, with more studies pointing towards a multifactorial process in which no single origin can be universally assumed (Hollund et al. 2012; Turner-Walker 2019; Turner-Walker et al. 2023; Schotsmans et al. 2024).

Cracking is one of the most frequently documented physical manifestations of bone diagenesis. It has been previously suggested that a freezing environment can increase bone cracking (Jans 2005; Kendall et al. 2018). However, the majority of available data derive from studies on faunal bones (Pelker et al. 1984; Letavernier and Ozouf 1987; Andrade et al. 2008; Fernández-Jalvo et al. 2010; Pfretzschner and Tütken 2011; Pokines et al. 2016; Turpin 2017), which raises the issue of limited applicability to human remains, given structural differences between human and animal bone (Turpin 2017; Oostraa et al. 2020; Schotsmans et al. 2024). Studies involving the analysis of humans were either focused on embalmed remains (Perkins 2012) or artificially frozen ones (Tersigni 2007). For instance, Tersigni (2007) noted the presence of Haversian cracks due to the freezing cycles artificially induced, while Trenchat and colleagues (under review) experimentally investigated the impact of a natural cold decomposition environment on modern human remains at a human taphonomy facility in Canada. Their preliminary data show alteration of bone microstructure by freeze-thaw cycles. However, the small sample size (N=11 human donors) and the comparatively short post-mortem interval in cold temperatures (10-36 months post-mortem) limit the generalisability of these results to archaeological contexts, where remains can be subject to centuries of climatic oscillations and long-term interaction with frozen soil.

Tunnug 1 is an archaeological and funerary site located in a frozen environment, more specifically situated in a swampy area on the river terraces of the Uyuk Valley (southern Siberia, Russia) (Figure 1). The region is known for its cold environment and its remarkable concentration of ancient funerary monuments, particularly large burial mounds and frozen tombs, which have been associated with the princely tombs of the Scythian culture and have been the subject of archaeological investigation since the 1970s (Caspari et al. 2018). These features posed significant challenges during excavations due to high groundwater levels, waterlogged conditions and extreme temperature variations between day and night (around 40°C of temperature variation between day and night) (Caspari et al. 2019). Tunnug 1 has been dated to the 9^th^ century BCE and represents one of the oldest known Scythian burial mounds (Caspari et al. 2018). The site showed evidence of use across multiple periods, ranging from the Late Bronze Age (15^th^-9^th^ centuries BCE) to the Turkic period (6^th^-8^th^ centuries CE) (Sadykov et al. 2020; Chan et al. 2022). Tunnug 1, like other funerary contexts of the Altai region, is characterised by frozen-soil or permafrost conditions, a factor that contributed to the exceptional preservation of the site. Permanently frozen soil (permafrost) was observed near the main burial mound (Belyaev et al. 2022), whereas funerary structures located in the southern periphery of the site were situated within an active frost layer, undergoing seasonally freezing and thawing (Caspari et al. 2018; Chan et al. 2022). Both continuous and discontinuous permafrost were documented during fieldwork, along with large frost cracks and evidence of cryo-bioturbations (Caspari et al. 2019). Consequently, the archaeological remains underwent environmental variation due to freeze-thaw cycles. Overall, these conditions make Tunnug 1 one of the largest examples of frozen tombs identified to date (Caspari et al. 2018).

**Figure 1:**
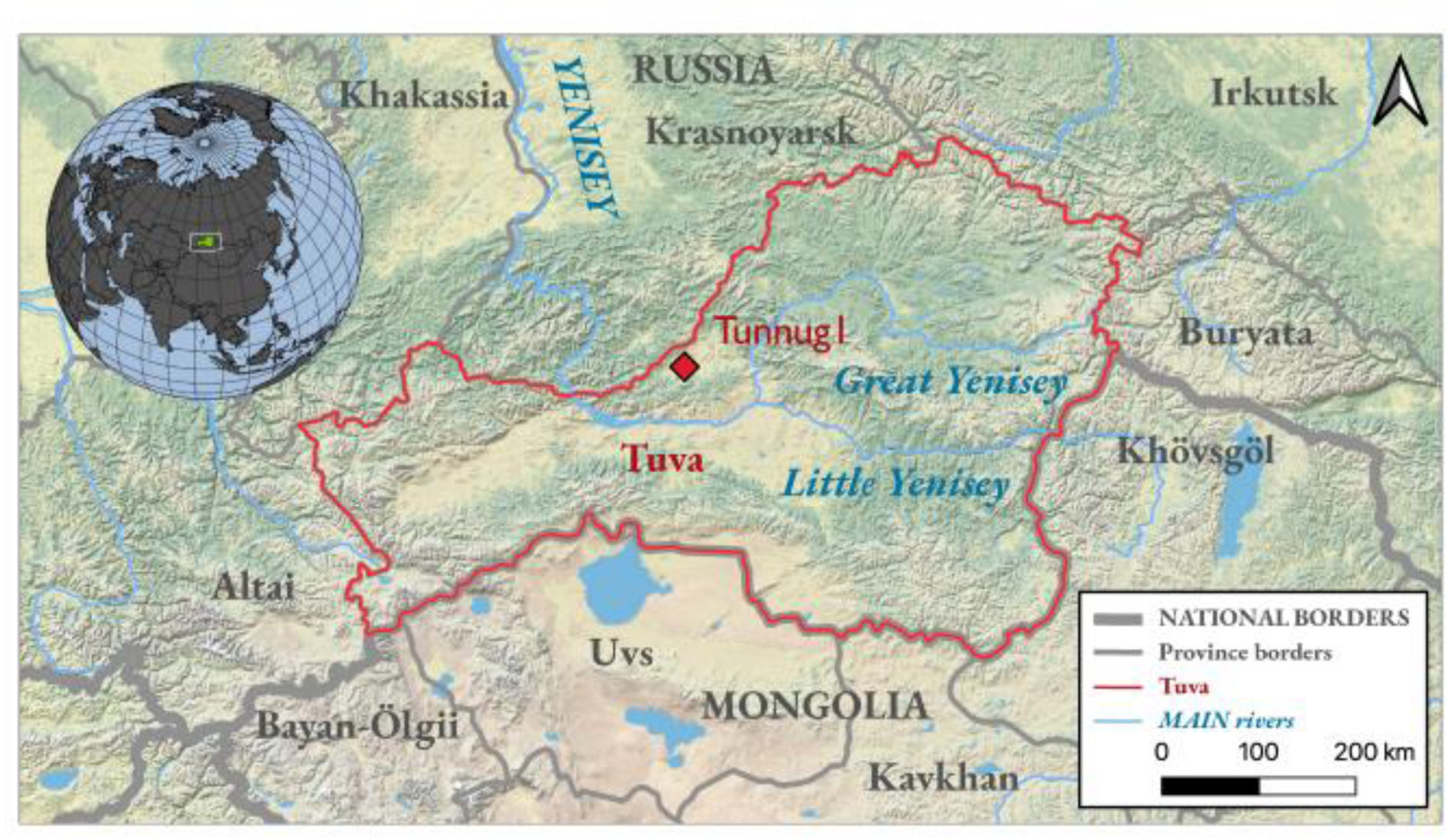
The location of Tunnug 1 in South Siberia (Milella et al. 2022).

Excavations at Tunnug 1 revealed a complex internal structure, organised with radial features and multiple chambers (Caspari et al. 2019). At the periphery of the mound, a cemetery associated with the Kokel culture (2^nd^-4^th^ centuries CE) was identified and excavated in 2018 and 2019 (Figure 2) (Caspari et al. 2018; Caspari et al. 2019). This indicates that the funerary use of the site was established after the construction of the mound itself. Funerary sites attributed to the Kokel culture are typically characterised by burial mounds of varying sizes, in which individuals were mostly buried in wooden coffins accompanied by grave goods, such as ceramics, knives, animal bone offerings and miniature objects (Sadykov et al. 2021).

**Figure 2:**
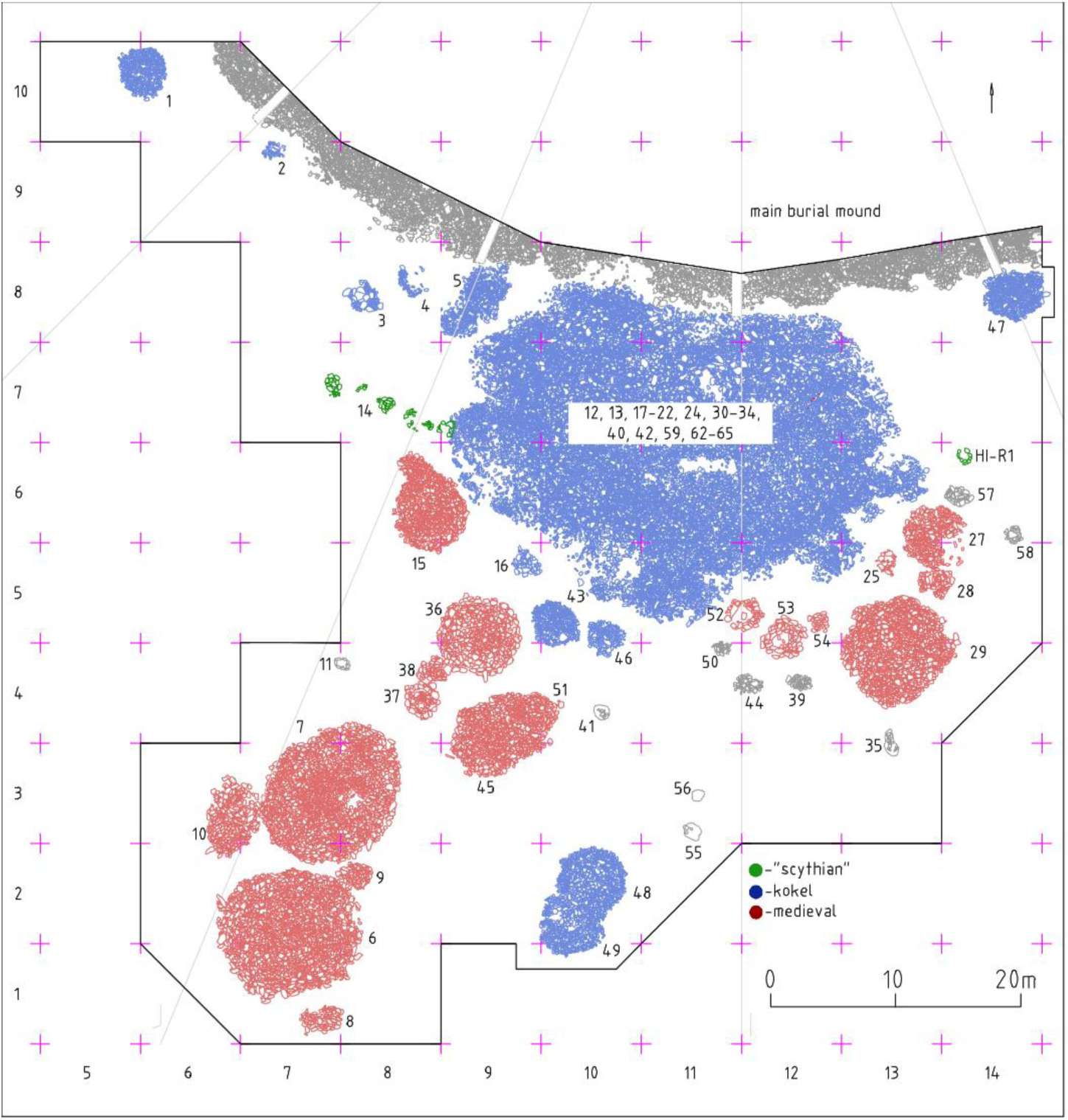
The Southern periphery of the burial mound of Tunnug 1, where the Kokel cemetery is located (Sadykov et al. 2021).

The funerary area of Tunnug 1 revealed principally burials from the Kokel period, some Scythian burials and medieval burials at the southern periphery (Sadykov et al. 2020) (Figure 2). In total, 46 inhumations were excavated: 39 single and 7 multiple burials with a minimum number of individuals of 87 individuals consisting of males and females from different age classes (Figure 3). Individuals were mainly placed in wooden coffins, in a supine and extended position, either in single pits under stone mounds or in multiple funerary structures containing individual coffins. Overall, the burials were well-preserved, with the exception of some inhumations disturbed by environmental factors, such as soil freezing and thawing, which induced soil frost cracks and frost-related substrate movement (Milella et al. 2020). The funerary material associated with Tunnug 1 was diverse: arrowheads, vessels, ceramics, wood, jewellery and gold objects (Sadykov et al. 2021). Similar grave goods were found in multiple and single graves (Milella et al. 2020; Sadykov et al. 2021; Sadykov et al. 2025). Isotopically (δ^34^S) individuals at Tunnug 1 appeared, with few exceptions, local (Milella et al. 2022), whereas the high prevalence of peri-mortem skeletal trauma and their distribution across sexes and age groups points to the exposure of these individuals to high levels of interpersonal violence (Milella et al. 2020), similarly to what was observed on skeletons from chronologically earlier contexts from the same region (Murphy 2003).

**Figure 3:**
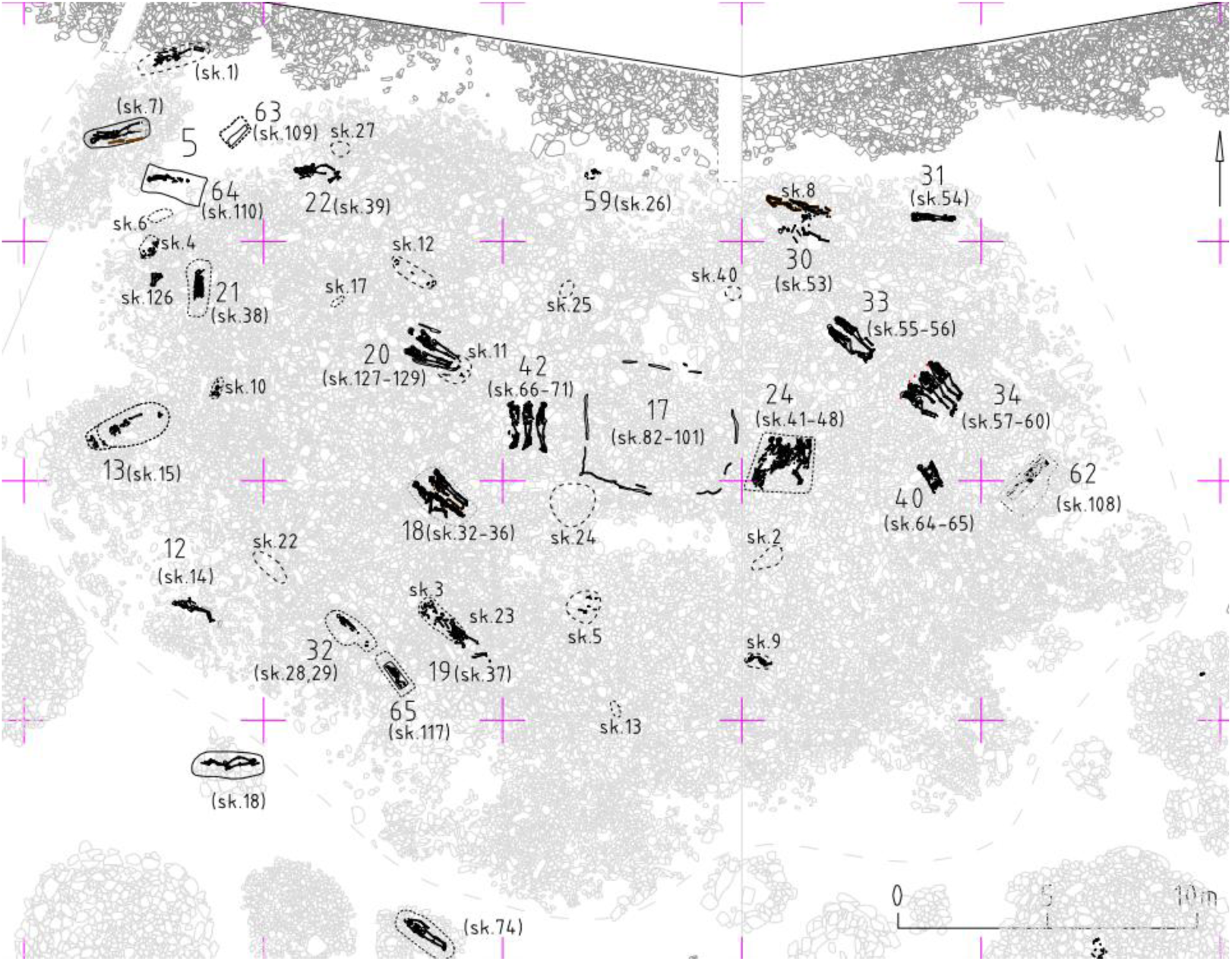
Tunnug 1: spatial distribution of burials (Sadykov et al. 2025).

Given this complex taphonomic history, assessing bone microstructure at Tunnug 1 requires methods capable of detecting both biological alteration and physical degradation, such as cracking. Bone degradation and cracking are often studied via transmitted light microscopy (resolution ≤ 1 μm) and Scanning Electron Microscopy (SEM). SEM is useful to identify MFD corresponding to chemical alterations represented by hyper- and de-mineralised areas. In addition, it allows observation of microcracks due to the highest resolution (approximately 0.5-4 nm) (Bell et al. 1991; Boyde and Jones 1996; Turner-Walker and Syversen 2002; Fernández-Jalvo et al. 2010; Dal Sasso et al. 2014; Vasquez et al. 2021; Turner-Walker et al. 2023; Schotsmans et al. 2024). Others use micro-computed tomography (micro-CT) to examine bone diagenesis in three-dimensions, but its low resolution (depending on the sample size, often between 5-15 μm for smaller samples) can limit the observation of diagenetic features (Booth et al. 2016; Caruso et al. 2020; Mandl et al. 2022; Loy et al. 2023; Trenchat et al. 2024). The most common scoring method for bone diagenesis is the Oxford Histological Index (OHI), developed by Hedges et al. (1995) and Millard (2001). This qualitative method aims to estimate the percentage of bone preservation based on histological observation and to score them from 0 (poorly preserved) to 5 (excellent preservation). Consequently, the resulting scores depend heavily on the quality of the thin section and are therefore affected by a degree of subjectivity (Schotsmans et al. 2024; Trenchat et al. 2024; Trenchat et al. 2025). To address this limitation, Trenchat et al. (2025) developed a quantitative approach to calculate a percentage of bone preservation using image analysis software. Rather than relying on subjective visual estimation, the method allows the isolation of preserved and degraded bone segments using density values represented through a greyscale histogram, enabling a reproducible percentage of preservation to be calculated (Trenchat et al. 2024; Trenchat et al. 2025).

Despite previous research on frozen environments, the long-term effects of repeated freeze-thaw cycles on bone preservation remain poorly understood, and it is unclear how funerary treatment interacts with these diagenetic processes. The Tunnug 1 skeletal assemblage offers a rare opportunity to address these questions: it spans individuals of all ages and sexes, buried across multiple periods and subject to diverse funerary treatments, including single and multiple burials, variable grave goods (i.e. iron objects) and the presence or absence of wooden coffins. Combining micro-CT and histological analysis on this assemblage allows both a comprehensive evaluation of bone preservation and a direct comparison of the strengths and limitations of each method. This study therefore aims to evaluate the preservation of bone microstructure at Tunnug 1, with the following objectives: (i) to assess bone preservation and characterise the diagenetic features present; (ii) to quantify the percentage of bone preservation and the percentage of cracking using image analysis; (iii) to statistically evaluate the relationship of cracking and bone degradation with funerary and environmental variables, including burial type (single or multiple), period, grave goods and the presence of coffins; (iv) and to examine variation in preservation at sample and individual level, including intra-individual variation and differences related to age (immature versus adult individuals).

## 2. Materials and methods

### 2.1. Materials

This study examined 74 human bone samples, either called Object (Obj) or Skeleton (Sk), from 36 burials, collected at Tunnug 1, altogether representing 71 individuals, as well as 2 faunal bone samples. The object number corresponds to the funerary structure observed and documented in the field. However, for some remains, the associated funerary structure could not be clearly identified and only a skeletal number was recorded. Samples were selected to represent the funerary variability of Tunnug 1 in terms of burial type (multiple and single burials), age categories and chronology (Kokel and Medieval period). Rib and long bone samples were collected, with one skeletal element per individual. For five individuals, both a rib and a long bone were sampled. For each individual, the presence or absence of a wooden coffin, grave goods (including iron objects) and trauma was recorded. Further details on burial context and anthropological data are presented in Supplementary Materials S1 and S2. More information can also be found in Sadykov et al. (2025).

Two structures, Objects 17 and 21, differed from the others. Object 17 corresponded to a mass burial that had been looted. It comprised a large wooden frame and 400 fragments of human bones belonging to at least 20 individuals (Milella et al. 2020; Sadykov et al. 2025). Structure 21 demonstrated a distinct funerary arrangement. The burial contained a well-preserved wooden coffin, comparable in length and structural characteristics to other coffins at the site and included similar grave goods such as ceramics. What set it apart, however, was the position of the individual: despite being placed in a coffin of standard and extended length, the body was arranged in a supine flexed position, unusual for the period (Sadykov et al. 2025).

### 2.2. Methods

#### 2.2.1. Biological anthropology

Prior to this study, an anthropological study was carried out on the skeletons from Tunnug 1 to determine their sex, age and paleopathology (Milella et al. 2020). Individuals were grouped in different age-at-death classes: neonates (up to 3 months at death), infants (4 months–3 years at death), children (3–12 years old at death), adolescents (13–18 years old), young adults (19–34 years old), middle adults (35–49 years old) and old adults (≥50 years old). For skeletons that were too fragmented, individuals were classified into broader groups: subadults (ca. <19 years old) and adults (ca. >19 years old). Further information on the methods used to determine the age and sex of the individuals is presented in Milella et al. (2020). A paleopathological and, specifically, paleotraumatological analysis was performed on the skeletons, given the high prevalence of peri-mortem trauma. Trauma was assessed and recorded by type, location and orientation. It included chop marks, slice marks, blunt trauma, penetrating lesions and fractures (Milella et al. 2020). In this study, the presence of trauma, grave goods and coffins was simply noted as present or absent.

#### 2.2.2. Micro-Computed Tomography (micro-CT)

Bone samples were scanned by micro-Computed Tomography at PACEA in Bordeaux (France). X-ray micro-CT scans were performed using a GE phoenix v|tome| x s scanner fitted with a 0.1 mm Cu filter. Every scan was conducted at 150 μA, 120 kV and 500 ms exposure. 2550 projections were acquired over a 360° rotation with 3 frames averaging. 3D images were reconstructed in 16 bits with the *Phoenix datos|x reconstruction 2* program. Depending on the bone geometry, the voxel size was set between 13 and 17 µm.

For each bone, the targeted scanned area was recorded to allow the thin section to be produced from the same anatomical location, as explained below. Once the 3D images were reconstructed, the slice matching the thin section was determined by comparing porosity patterns and morphological features visible both in optical microscopy and microtomography rendering. This slice is referred to as the ‘corresponding slice’ throughout this research.

Micro-CT scanning could not be performed on Skeleton 4, Skeleton 6, Object 24_skeleton 6 and the faunal bones, as these samples were not available at the time of acquisition; these individuals are therefore absent from the microtomographic dataset.

#### 2.2.3. Histological protocol

Every sample was processed into thin sections at the University of Wollongong (Australia). For each sample, the thin section was located within the volume previously scanned by microtomography. Standard histological methods were followed to produce the thin sections (Bancroft and Gamble 2002; Miszkiewicz and Mahoney 2017; Miszkiewicz 2019).

The bone samples were embedded in Buehler Epothin epoxy resin (2 to 1 ratio of resin and hardener) in plastic moulds with a removable lid (Telfon moulds of 25mm). A mould release spray (Silfree-jet aerosol) was applied to facilitate demoulding of the embedded samples. Each embedded block was polished using a motorised rotary grinder-polisher (Presi, Le Cube), progressing through four grinding discs of decreasing grit (P240, P220, P600 and P1200 SiC paper), followed by polishing with diamond suspensions (6 µm, 3 µm and 1 µm) on Kan-M and ASFL-M cloths. The polished blocks were then mounted on glass slides (76 × 26 mm) using the same epoxy resin as used for embedding. Sections were cut with a precision diamond-bladed saw (Buehler Isomet 1000; blade: 152 × 0.5 mm) and subsequently polished following the same sequence of steps until a final thickness between 80–200 µm was achieved.

#### 2.2.4. Microscopic analysis

The thin sections were examined under transmitted and polarised light using an Olympus BX51 microscope equipped with a DP23 high-resolution camera and operated by cellSens Olympus software. Each thin section was fully auto-stitched at x100 total magnification using the cellSens instant microscope image analysis tool. In addition, higher magnification images (x200 and x400) were acquired for numerical resolution between 1 and 4 μm/pixel and a field of view of 1525 x 808 pixels.

The embedded samples were further analysed using a Phenom XL Scanning Electron Microscope (SEM) equipped with a silicon drift detector in low-vacuum mode. Backscattered electron images were acquired at an accelerating voltage of 15 kV and elemental analyses were performed using Energy-dispersive X-Ray spectroscopy (EDX) with PhenomProSuite software. In backscattered electron images, contrast is determined by variations in atomic number density. Bone degradation is characterised by hyper-mineralised areas, which appear bright due to their higher density and mineral content, and demineralised areas, which appear darker as a result of reduced density and mineral content (Turner-Walker and Syversen 2002). Entire bone sections were mapped at low magnification (depending on the sample size and with numerical resolution around 1 μm/pixel) and detailed images were taken at higher magnifications x350 and x500 (numerical resolution between 0.25 and 0.35 μm/pixel and field of view of 2048 x 2176 pixels).

#### 2.2.5. Assessment of bone degradation

Bone diagenesis was assessed using the OHI, ranging from 0 to 5 (0: 0–5% poorly preserved; 1: 5–15% bone matrix is destroyed by the presence of MFD; 2: 15–50% some well-preserved bone is identified between the destroyed bone areas; 3: 50–85% large parts of well-preserved bone; 4: 85–95% bone well-preserved; 5: 95–100% excellent bone preservation) (Hedges et al. 1995; Millard 2001).

For each sample, bone diagenesis has been assessed through three complementary analyses: the entire micro-CT scan, the corresponding slice and the thin section observed under transmitted light microscopy. For each modality, the traditional OHI was scored twice by the same observer at two separate time points to assess intra-observer consistency. This scoring is referred to as the ‘Traditional OHI’ throughout the rest of the paper. In addition, bone preservation was quantified for each analysis using the image analysis method developed by Trenchat et al. (2025), which segments preserved from degraded bone on the basis of greyscale density values using Otsu’s thresholding algorithm (Figure 4). The segmentation was carried out with Dragonfly (Comet Technologies Canada Inc. (2025)). The resulting scores are referred to as the ‘Quantitative OHI’. All scores and percentages are provided in the Supplementary Material (S3) and were subjected to statistical testing.

**Figure 4:**
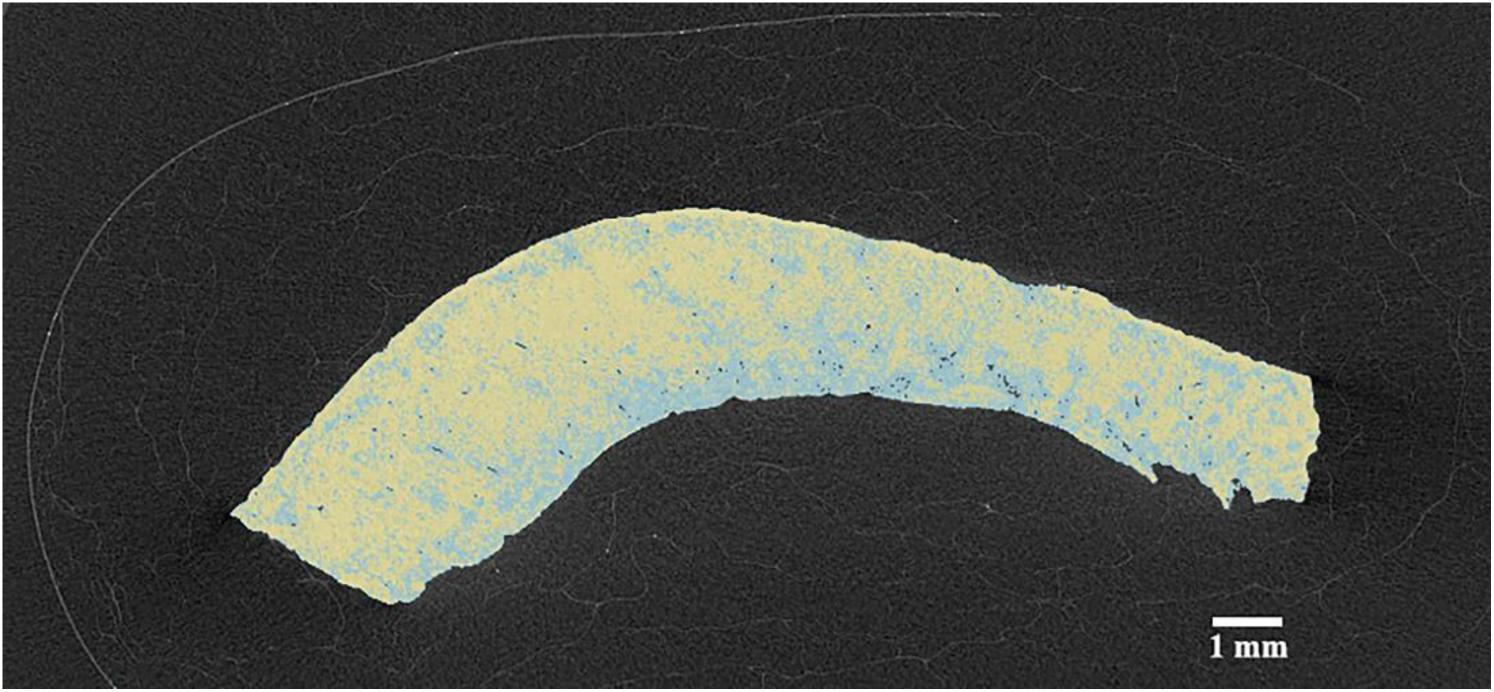
Micro-CT scan of Skeleton 17 (tibia) with preserved bone segmented (in yellow) from altered bone (in blue) to allow bone preservation quantification.

#### 2.2.6. Assessing bone cracking

The same segmentation method, using the Otsu algorithm in Dragonfly, was applied to quantify the percentage of cracking and microcracking on the generated SEM overview images. The higher resolution of SEM allows for the detection of finer microcracks that would otherwise remain invisible (Schotsmans et al. 2024). The cracks were first isolated from the rest of the bone structure by segmentation. Then a cracking percentage was calculated in relation to the whole bone area. Since bone contains natural pores, such as Haversian canals, these features were separated from actual cracks by the software. However, in most cases, some small pores had to be removed manually, using ‘Remove island’ tools in the software. By setting thresholds, large cracks (considered as overall cracks, approximately <5 μm) were distinguished from microcracks (approximately > 5 μm).

#### 2.2.7. Statistical analysis

Bone preservation was assessed using quantified continuous percentages. Bone cracking, corresponding to the areas of cracks, was also examined using continuous percentages. The sex of samples could not be tested as many individuals had an indeterminate sex and could not be included in the statistical analysis. The iron objects and the coffins were noted as present or absent. The individuals for the period ‘likely Kokel’ were considered from the Kokel period for the analysis. Age class categories were combined into three broad categories to increase the sample size of each level: Immature (including neonates, infant and child categories), Subadult (including adolescent and subadult) and Adult (including young adult, middle adult, adult and old adult). Bone type was classed into two general categories: Long bone and Rib. Burial depth was recorded in metres.

Preliminary data exploration showed strong associations among several explanatory variables and incomplete representation of some combinations of predictor values. For example, the deeper burials were exclusively associated with coffin use, iron objects occurred only in coffin burials, and long bone samples were disproportionately represented at the greatest burial depths. These relationships limited the ability to estimate independent effects reliably within a multivariable framework. For this reason, analyses focused on univariate comparisons between each explanatory variable and the response variables. Non-parametric tests (Mann-Whitney U and Kruskal-Wallis) were used because of unequal group sizes and did not consistently satisfy assumptions of normality. Associations between the response variables and burial depth were assessed using Spearman’s rank correlation coefficient.

All analyses were conducted in R (Team 2025). Statistical significance was evaluated at an alpha of 0.05. Statistical details, including the R code and the data, are provided in the Supplementary Material (S4).

## 3. Results

### 3.1. Bone degradation: OHI scores

The traditional OHI of the thin sections and the quantitative OHI from the micro-CT scans are presented in Table 1. The bone samples from Tunnug 1 presented a bone preservation ranging between OHI 2 and 4. Differences between the traditional OHI and the quantitative OHI were noted for 33 samples. The majority was observed between two consecutive categories, most frequently between scores 3 and 4. In contrast, only three samples (Object 5, Object 24_skeleton 3, Object 42_skeleton 5) exhibited differences between non-consecutive categories (scores 2 and 4).

**Table 1:**
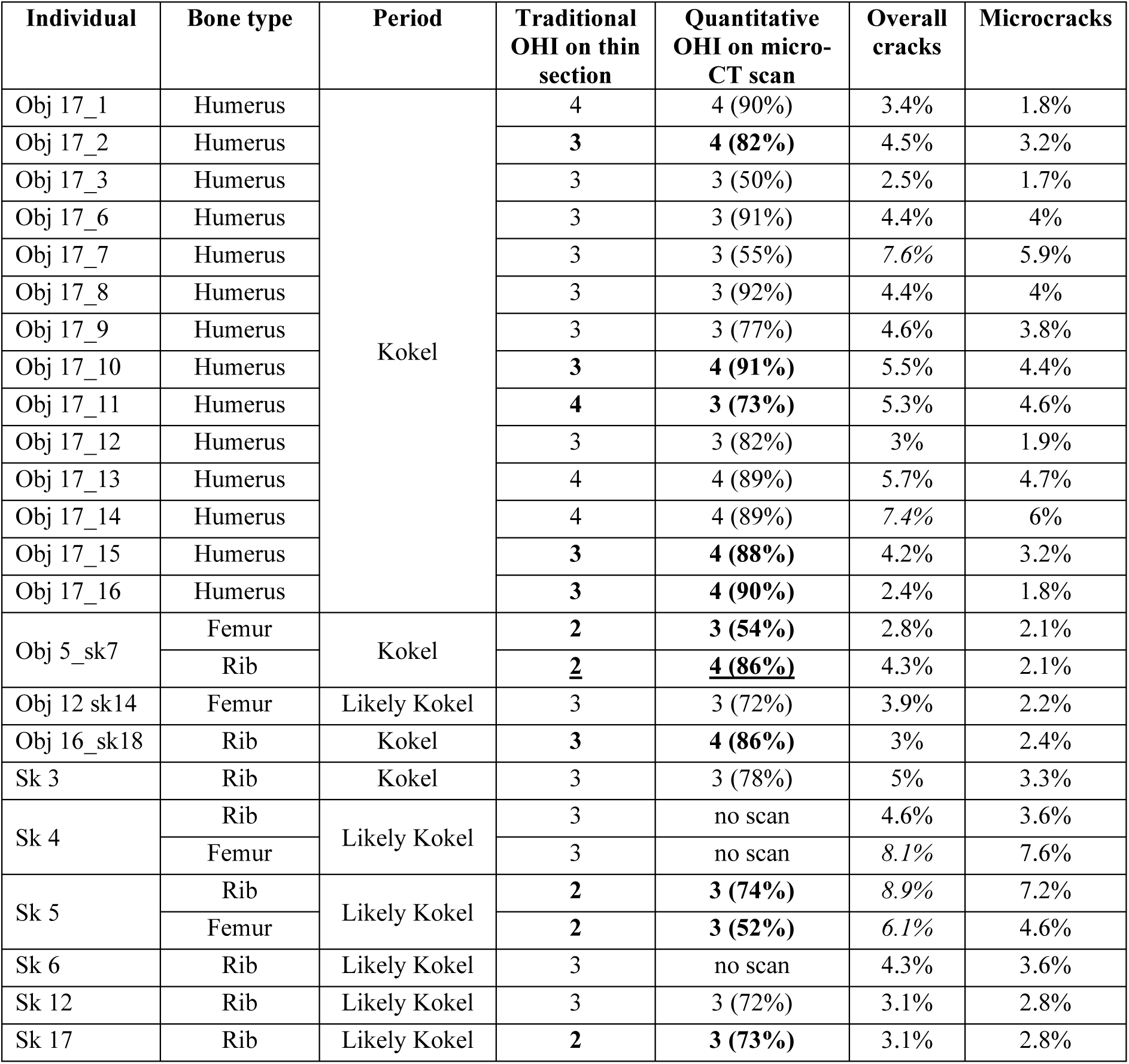
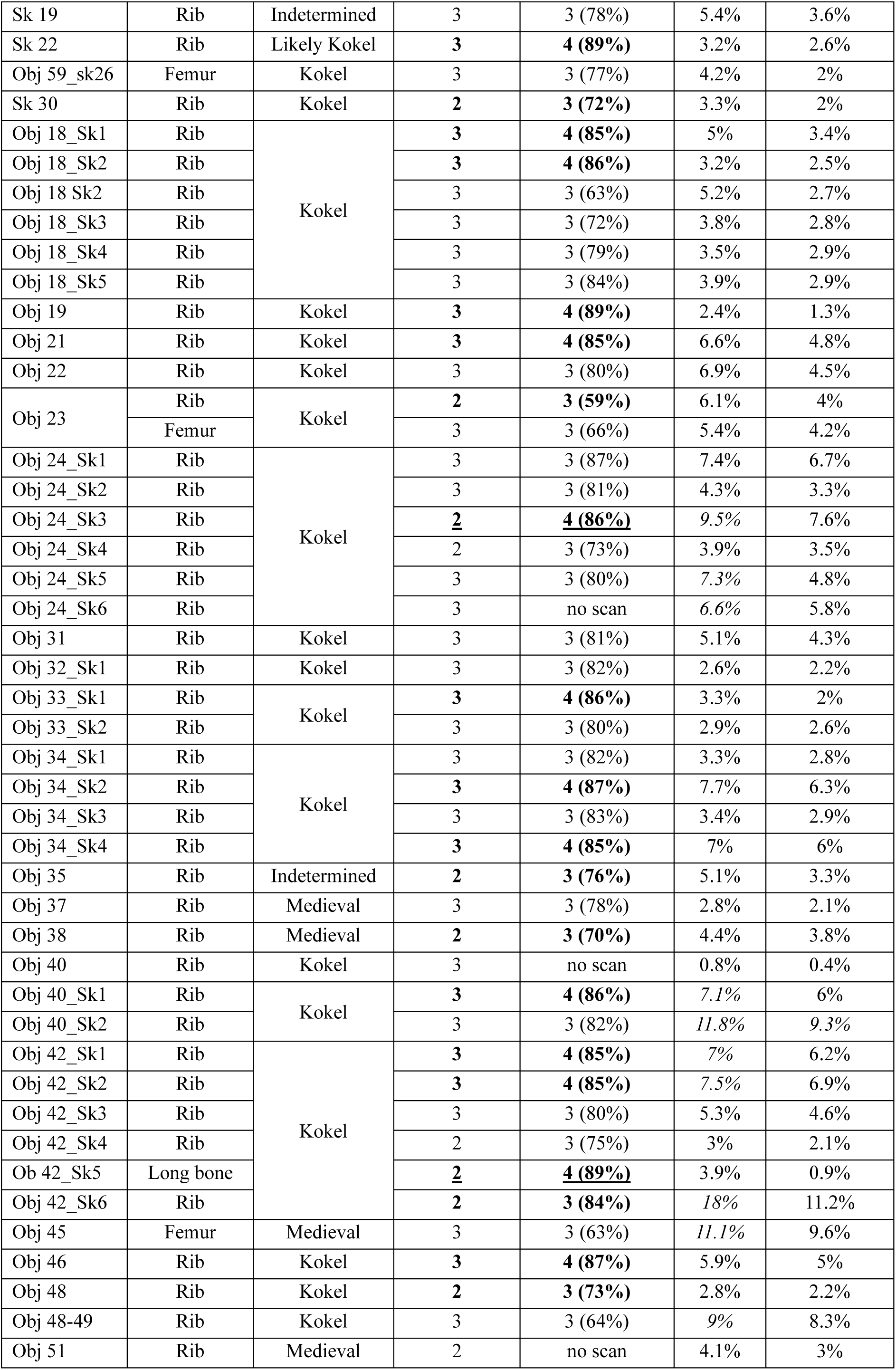
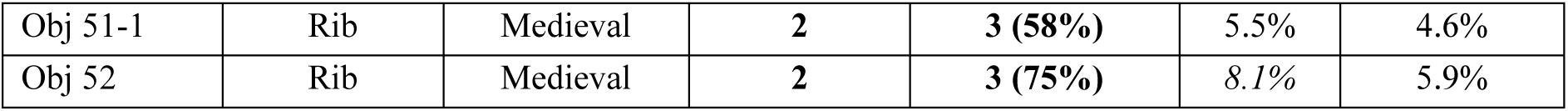
Bone type and period time from each sample. Traditional OHI of thin sections and quantitative OHI on the micro-CT scans. In bold: difference between two consecutive categories of scores of the same sample; underlined: differences between two non-consecutive categories. Percentages of cracks and microcracks, in italic percentages >5%.

| Individual | Bone type | Period | Traditional OHI on thin section | Quantitative OHI on micro-CT scan | Overall cracks | Microcracks |
| --- | --- | --- | --- | --- | --- | --- |
| Obj 17_1 | Humerus | Kokel | 4 | 4 (90%) | 3.4% | 1.8% |
| Obj 17_2 | Humerus |  | **3** | **4 (82%)** | 4.5% | 3.2% |
| Obj 17_3 | Humerus |  | 3 | 3 (50%) | 2.5% | 1.7% |
| Obj 17_6 | Humerus |  | 3 | 3 (91%) | 4.4% | 4% |
| Obj 17_7 | Humerus |  | 3 | 3 (55%) | 7.6% | 5.9% |
| Obj 17_8 | Humerus |  | 3 | 3 (92%) | 4.4% | 4% |
| Obj 17_9 | Humerus |  | 3 | 3 (77%) | 4.6% | 3.8% |
| Obj 17_10 | Humerus |  | **3** | **4 (91%)** | 5.5% | 4.4% |
| Obj 17_11 | Humerus |  | **4** | **3 (73%)** | 5.3% | 4.6% |
| Obj 17_12 | Humerus |  | 3 | 3 (82%) | 3% | 1.9% |
| Obj 17_13 | Humerus |  | 4 | 4 (89%) | 5.7% | 4.7% |
| Obj 17_14 | Humerus |  | 4 | 4 (89%) | 7.4% | 6% |
| Obj 17_15 | Humerus |  | **3** | **4 (88%)** | 4.2% | 3.2% |
| Obj 17_16 | Humerus |  | **3** | **4 (90%)** | 2.4% | 1.8% |
| Obj 5_sk7 | Femur | Kokel | **2** | **3 (54%)** | 2.8% | 2.1% |
|  | Rib |  | **<u>2</u>** | **<u>4 (86%)</u>** | 4.3% | 2.1% |
| Obj 12 sk14 | Femur | Likely Kokel | 3 | 3 (72%) | 3.9% | 2.2% |
| Obj 16_sk18 | Rib | Kokel | **3** | **4 (86%)** | 3% | 2.4% |
| Sk 3 | Rib | Kokel | 3 | 3 (78%) | 5% | 3.3% |
| Sk 4 | Rib | Likely Kokel | 3 | no scan | 4.6% | 3.6% |
|  | Femur |  | 3 | no scan | 8.1% | 7.6% |
| Sk 5 | Rib | Likely Kokel | **2** | **3 (74%)** | 8.9% | 7.2% |
|  | Femur |  | **2** | **3 (52%)** | 6.1% | 4.6% |
| Sk 6 | Rib | Likely Kokel | 3 | no scan | 4.3% | 3.6% |
| Sk 12 | Rib | Likely Kokel | 3 | 3 (72%) | 3.1% | 2.8% |
| Sk 17 | Rib | Likely Kokel | **2** | **3 (73%)** | 3.1% | 2.8% |
| Sk 19 | Rib | Indetermined | 3 | 3 (78%) | 5.4% | 3.6% |
| Sk 22 | Rib | Likely Kokel | **3** | **4 (89%)** | 3.2% | 2.6% |
| Obj 59_sk26 | Femur | Kokel | 3 | 3 (77%) | 4.2% | 2% |
| Sk 30 | Rib | Kokel | **2** | **3 (72%)** | 3.3% | 2% |
| Obj 18_Sk1 | Rib | Kokel | **3** | **4 (85%)** | 5% | 3.4% |
| Obj 18_Sk2 | Rib |  | **3** | **4 (86%)** | 3.2% | 2.5% |
| Obj 18_Sk2 | Rib |  | 3 | 3 (63%) | 5.2% | 2.7% |
| Obj 18_Sk3 | Rib |  | 3 | 3 (72%) | 3.8% | 2.8% |
| Obj 18_Sk4 | Rib |  | 3 | 3 (79%) | 3.5% | 2.9% |
| Obj 18_Sk5 | Rib |  | 3 | 3 (84%) | 3.9% | 2.9% |
| Obj 19 | Rib | Kokel | **3** | **4 (89%)** | 2.4% | 1.3% |
| Obj 21 | Rib | Kokel | **3** | **4 (85%)** | 6.6% | 4.8% |
| Obj 22 | Rib | Kokel | 3 | 3 (80%) | 6.9% | 4.5% |
| Obj 23 | Rib | Kokel | **2** | **3 (59%)** | 6.1% | 4% |
|  | Femur |  | 3 | 3 (66%) | 5.4% | 4.2% |
| Obj 24_Sk1 | Rib | Kokel | 3 | 3 (87%) | 7.4% | 6.7% |
| Obj 24_Sk2 | Rib |  | 3 | 3 (81%) | 4.3% | 3.3% |
| Obj 24_Sk3 | Rib |  | **2** | **4 (86%)** | 9.5% | 7.6% |
| Obj 24_Sk4 | Rib |  | 2 | 3 (73%) | 3.9% | 3.5% |
| Obj 24_Sk5 | Rib |  | 3 | 3 (80%) | 7.3% | 4.8% |
| Obj 24_Sk6 | Rib |  | 3 | no scan | 6.6% | 5.8% |
| Obj 31 | Rib | Kokel | 3 | 3 (81%) | 5.1% | 4.3% |
| Obj 32_Sk1 | Rib | Kokel | 3 | 3 (82%) | 2.6% | 2.2% |
| Obj 33_Sk1 | Rib | Kokel | **3** | **4 (86%)** | 3.3% | 2% |
| Obj 33_Sk2 | Rib |  | 3 | 3 (80%) | 2.9% | 2.6% |
| Obj 34_Sk1 | Rib | Kokel | 3 | 3 (82%) | 3.3% | 2.8% |
| Obj 34_Sk2 | Rib |  | **3** | **4 (87%)** | 7.7% | 6.3% |
| Obj 34_Sk3 | Rib |  | 3 | 3 (83%) | 3.4% | 2.9% |
| Obj 34_Sk4 | Rib |  | **3** | **4 (85%)** | 7% | 6% |
| Obj 35 | Rib | Indetermined | **2** | **3 (76%)** | 5.1% | 3.3% |
| Obj 37 | Rib | Medieval | 3 | 3 (78%) | 2.8% | 2.1% |
| Obj 38 | Rib | Medieval | **2** | **3 (70%)** | 4.4% | 3.8% |
| Obj 40 | Rib | Kokel | 3 | no scan | 0.8% | 0.4% |
| Obj 40_Sk1 | Rib | Kokel | **3** | **4 (86%)** | 7.1% | 6% |
| Obj 40_Sk2 | Rib |  | 3 | 3 (82%) | 11.8% | 9.3% |
| Obj 42_Sk1 | Rib | Kokel | **3** | **4 (85%)** | 7% | 6.2% |
| Obj 42_Sk2 | Rib |  | **3** | **4 (85%)** | 7.5% | 6.9% |
| Obj 42_Sk3 | Rib |  | 3 | 3 (80%) | 5.3% | 4.6% |
| Obj 42_Sk4 | Rib |  | 2 | 3 (75%) | 3% | 2.1% |
| Obj 42_Sk5 | Long bone |  | **2** | **4 (89%)** | 3.9% | 0.9% |
| Obj 42_Sk6 | Rib |  | **2** | **3 (84%)** | 18% | 11.2% |
| Obj 45 | Femur | Medieval | 3 | 3 (63%) | 11.1% | 9.6% |
| Obj 46 | Rib | Kokel | **3** | **4 (87%)** | 5.9% | 5% |
| Obj 48 | Rib | Kokel | **2** | **3 (73%)** | 2.8% | 2.2% |
| Obj 48-49 | Rib | Kokel | 3 | 3 (64%) | 9% | 8.3% |
| Obj 51 | Rib | Medieval | 2 | no scan | 4.1% | 3% |
| Obj 51-1 | Rib | Medieval | **2** | **3 (58%)** | 5.5% | 4.6% |
| Obj 52 | Rib | Medieval | **2** | **3 (75%)** | 8.1% | 5.9% |

### 3.2. Bone diagenesis: features

Three individuals presented biological degradation: Skeleton 5, Skeleton 12 and Skeleton 17, which were observable in the three analytic methods (micro-CT scan, transmitted light microscopy and SEM), excluding the possibility that they resulted from histological preparation (Figures 5 and 6). MFD was represented by less dense areas on the micro-CT scans, by black spots on the histological sections observed with a transmitted light microscope (Figure 5a) and by hyper-and de-mineralised areas on SEM images (Figure 5b). However, the degree of degradation varied considerably between individuals: Skeleton 5 exhibited the most severe deterioration (52%) and an overall cracking percentage (18%), suggesting that biological degradation had substantially weakened the bone matrix and promoted crack formation. Skeletons 12 and 17, by contrast, showed more limited areas of MFD (Figure 5).

**Figure 5:**
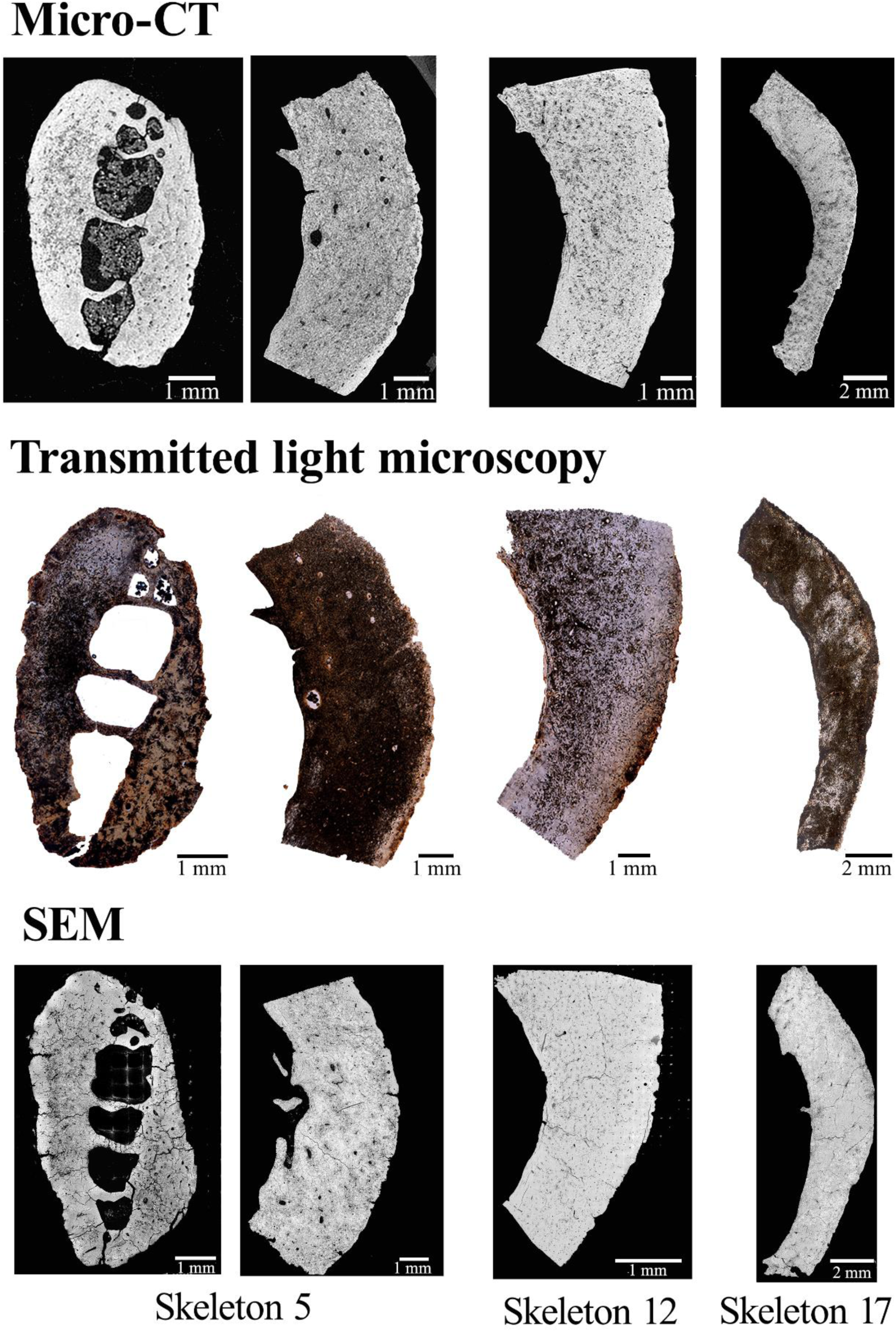
Rib and femoral bone samples from Skeletons 5, femoral bone sample from Skeleton 12 and tibia bone sample from Skeleton 17, all observed with micro-CT scan, transmitted light microscopy and SEM, demonstrating MFD in the bone structures: darker areas for the micro-CT scan, black areas for transmitted light microscopy and darker areas for SEM images.

**Figure 6:**
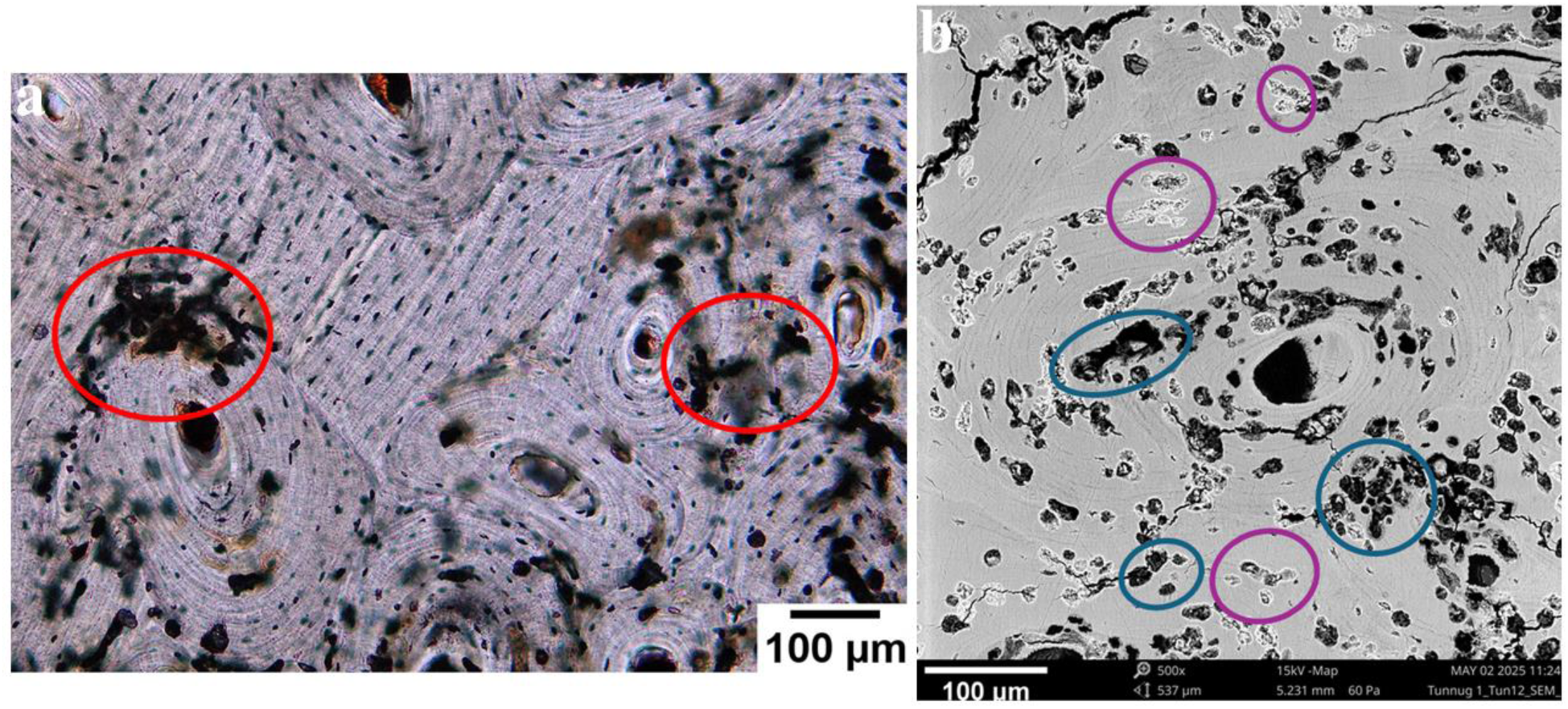
Detail of MFD from the femoral sample from Skeleton 12. 6a. Transmitted light microscopy: MFD correspond to black spots (red circles). 6b. SEM: MFD correspond to demineralised (blue circles) and hyper-mineralised (purple circles) areas.

Skeleton 30 exhibited localised degradation of the rib, characterised by rounded holes visible in micro-CT and histological images (Figure 7). These holes were confined to a specific area of the pleural surface. Suggesting a localised rather than diffuse diagenetic process.

**Figure 7:**
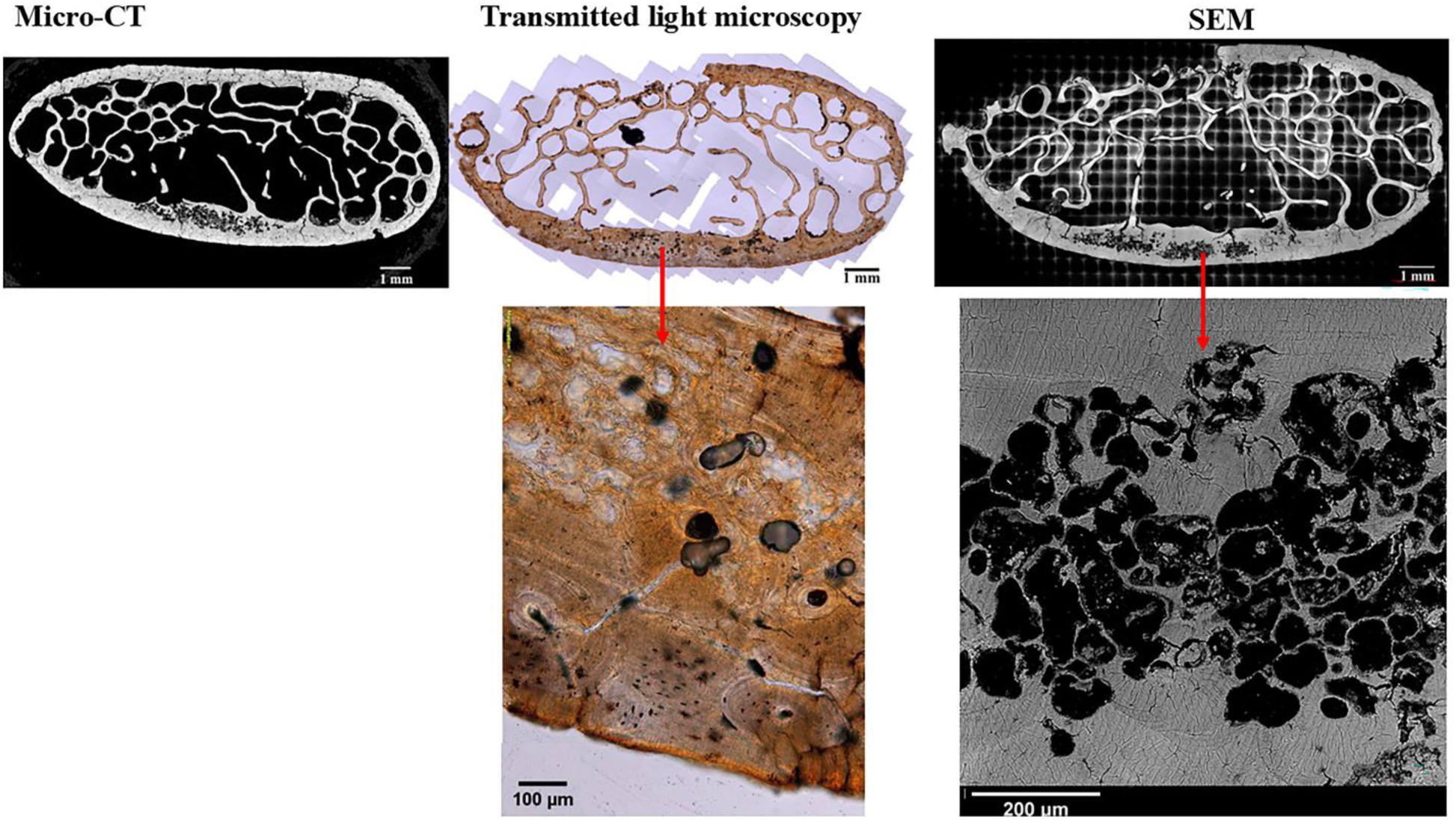
Chemical degradation observed on the rib of Skeleton 30, observed with micro-CT scan, transmitted light microscopy and SEM.

The rib from Object 23 displayed rounded holes in the bone structure under transmitted light microscopy similar to the ones observed on Skeleton 30 (Figure 8). The section also exhibited marked bone porosity, reduced bone density and demineralised areas. SEM-EDX analysis revealed variations in calcium and phosphorus ratios, as illustrated in Figure 8, where three distinct zones from the rib of Object 23 were examined. Hyper-mineralised bone (point 3 in Figure 8) exhibited higher calcium and phosphate ratios than demineralised bone (point 2). However, both altered areas displayed lower ratios than the unaltered bone (point 1). Extensive soil infiltration was additionally noticed, which reduced the visibility of the bone microstructure. The neonate individual from Object 51-1 similarly presented holes in the bone structure, soil infiltration and demineralised areas (Figure 9).

**Figure 8:**
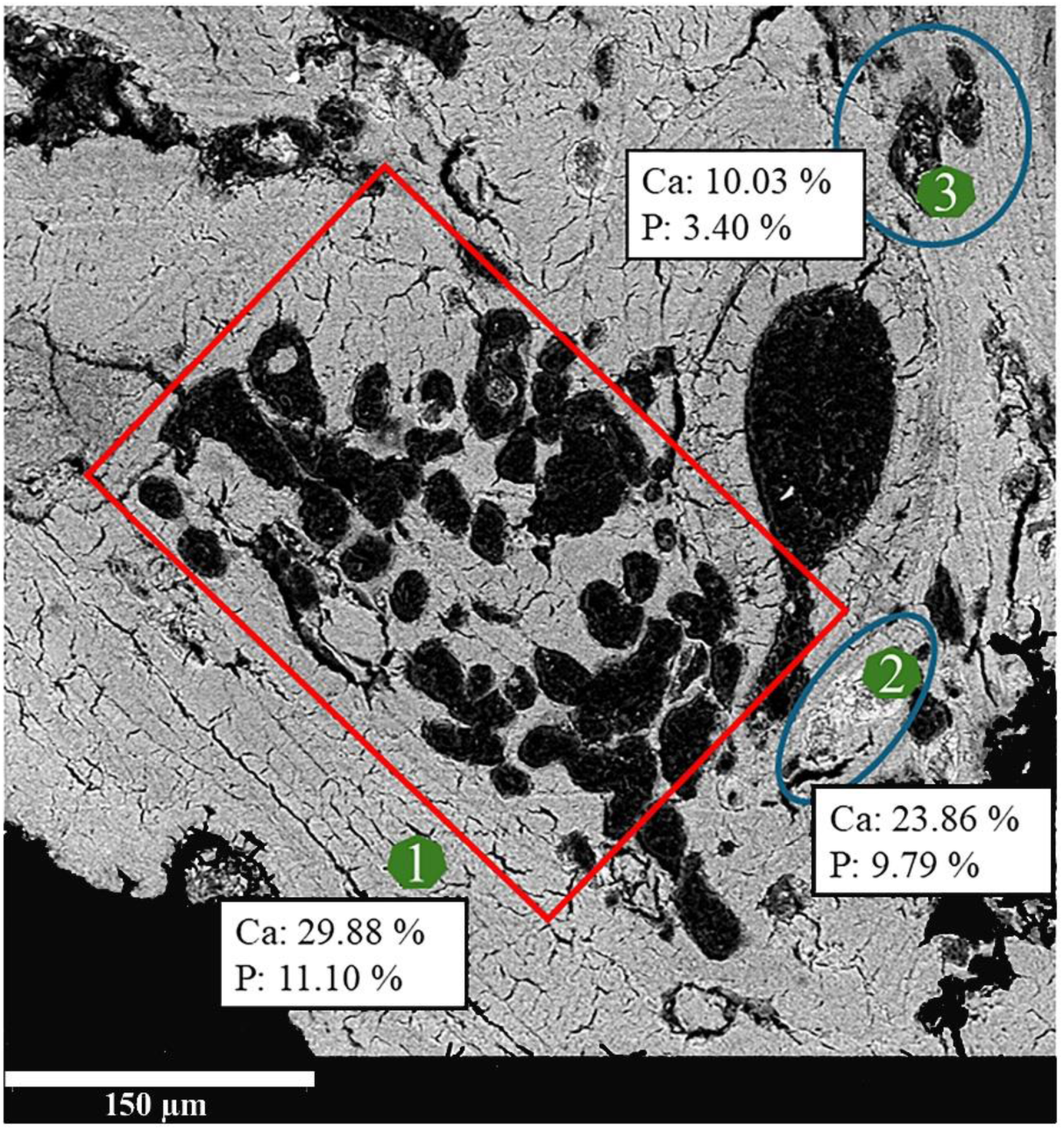
Rib from Object 23 presenting holes in the bone structure (within red square), de- and hyper-mineralised areas (blue circles). Elemental results with the weight percentages on three different locations (in green): point 1 corresponding to unaltered bone, point 2 corresponding to hyper-mineralised bone and point 3 to demineralised bone (Ca: Calcium, P: Phosphate).

**Figure 9:**
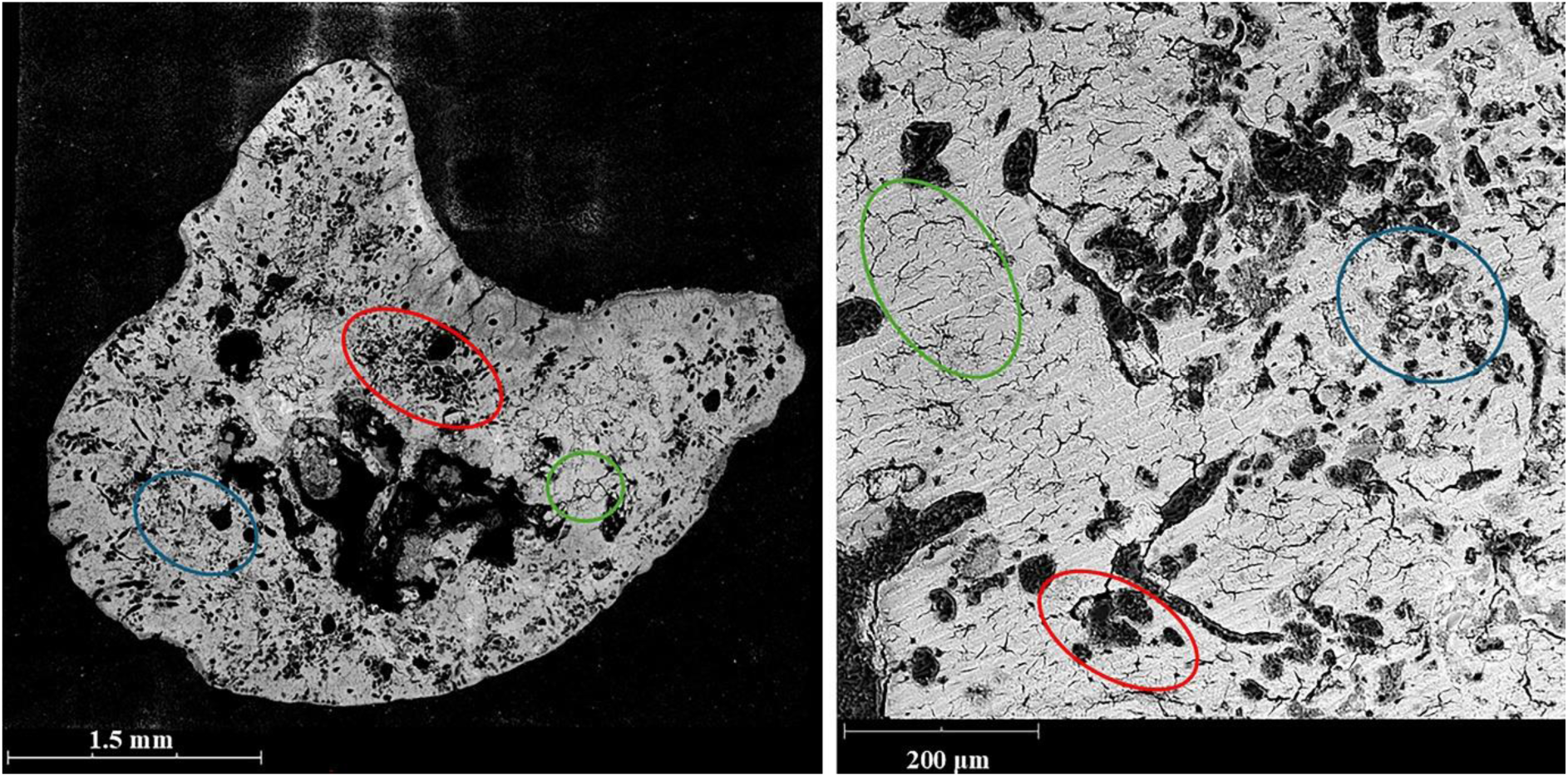
Object 51-1 (Rib sample, Neonate) observed with SEM (overview and detail images), presenting de- and hyper-mineralised areas (blue circles), bone cracking (green circles) and holes in the bone structure (red circles).

Enlarged canaliculi were observed on every bone sample with transmitted light microscopy (Figure 10). All bone samples displayed staining on their external parts, affecting both periosteal and endosteal surfaces of the cortical long bones and the pleural and cutaneous surfaces of the ribs. With a transmitted light microscope, staining was represented by an orange discolouration, while on SEM and microtomographic images, they were observed by whiter areas (Figure 5). SEM-EDX analysis identified the staining as originating from soil infiltration, evidenced by the presence of silicon and aluminium, as well as iron. The stained areas appeared less degraded than the rest of the bones. For example, Skeleton 5’s femoral sample appeared completely attacked by tunnels of degradation except for the periosteum part, where iron infiltrations were observed (Figure 5).

**Figure 10:**
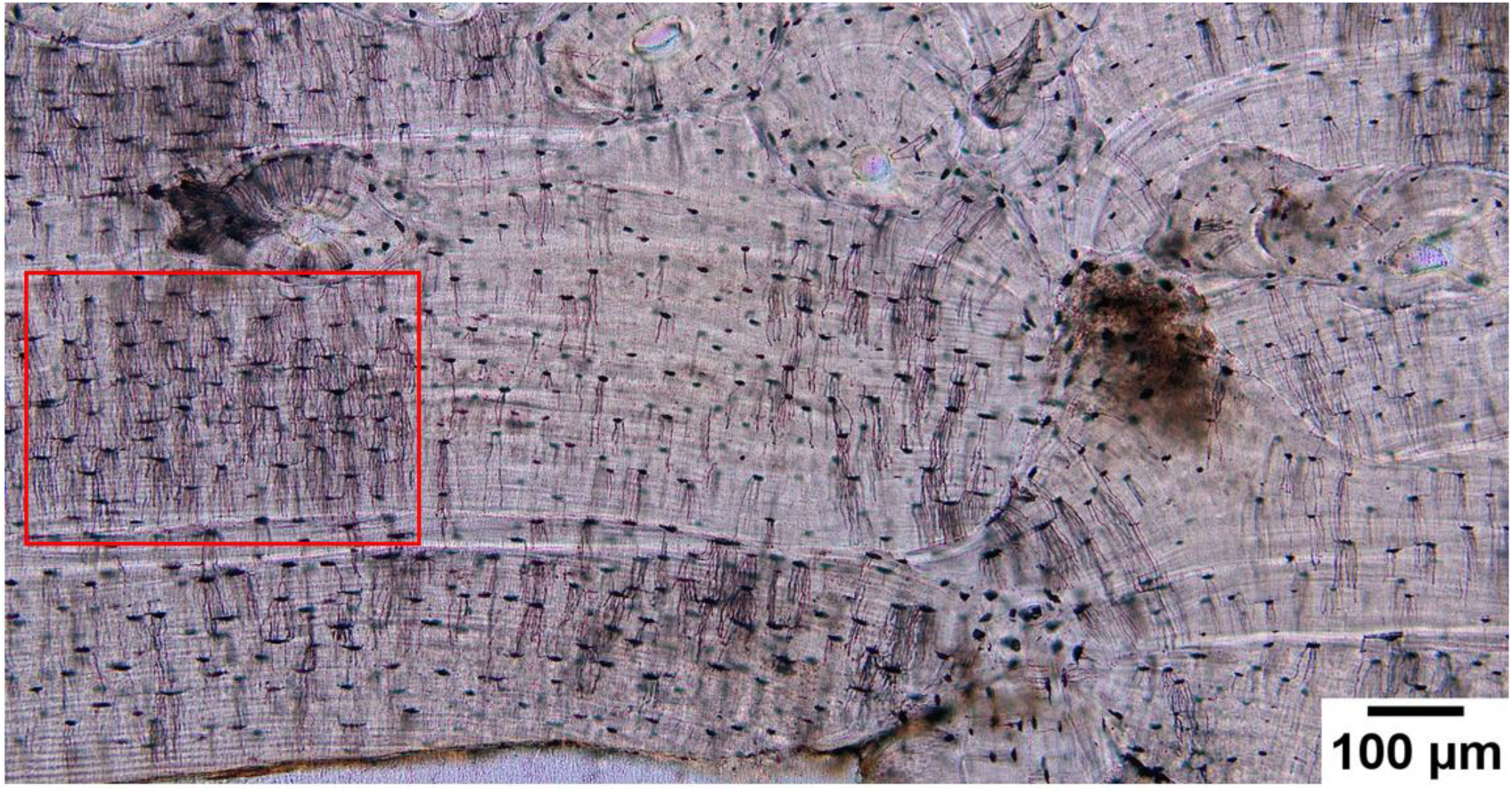
Object 19 observed under transmitted light microscopy, revealing enlarged canaliculi in the bone structure (red square).

### 3.4. Variations between samples and individuals

#### 3.4.1. Intra-individual variation

Two individuals were assessed on intra-individual bone preservation: Skeleton 5 and Object 23, both of which had a rib and a femur sampled. In Skeleton 5, both elements displayed comparable levels of degradation. Object 23, however, showed marked variation, with the rib exhibiting substantially greater degradation and cracking than the femur from the same individual (Table 1).

#### 3.4.2. Immature individuals

The neonate individuals (Object 35, Object 48 and Object 51-1) demonstrated poor bone preservation (OHI 2-3) with significant soil infiltration, pronounced bone porosity and an orange discolouration visible under transmitted light microscopy (Figure 11).

**Figure 11:**
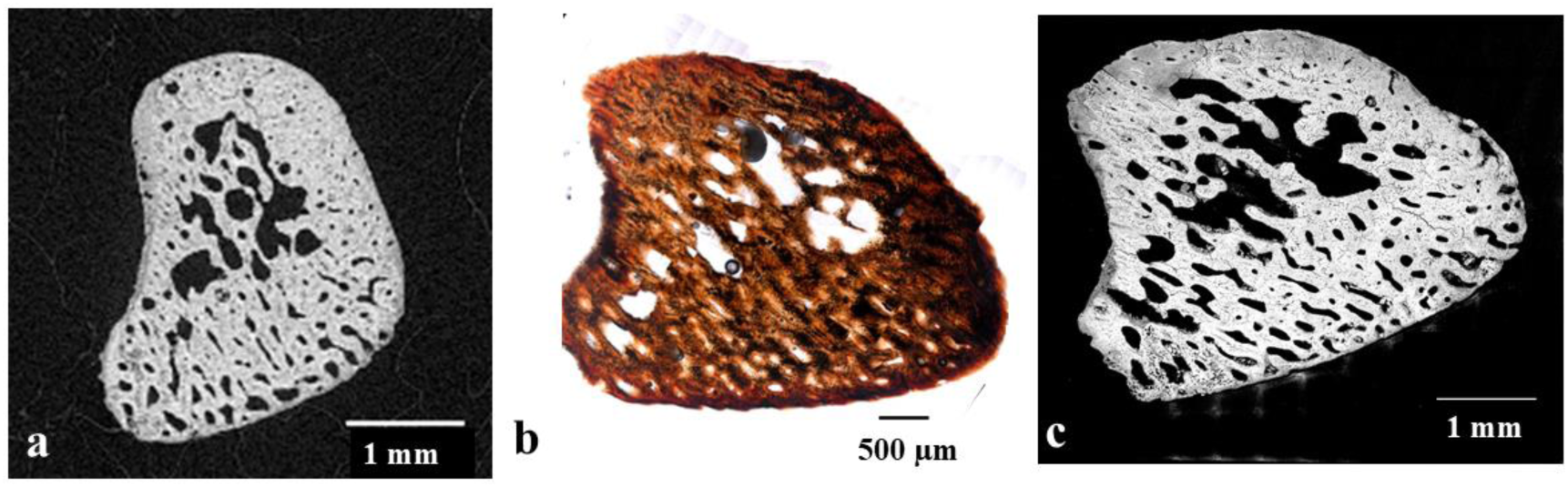
Object 48 observed with micro-CT scan (a), transmitted light microscopy (b) and SEM (c).

### 3.5. Cracking

Microcracks were identified in every sample, exclusively visible with SEM. The microcracks were located both within the osteonal structure and in the interstitial areas between osteons (Figure 12). The percentage of overall cracking and microcracking was quantified for each sample using the image analysis method in Dragonfly, with results presented in Table 1. Overall cracking percentages ranged from 1% to 18% across the assemblage. The most heavily cracked samples corresponded to those exhibiting the greatest degree of bone degradation, notably Skeleton 5 (18% and 11.1%), Object 51-1 (11.8%) and Skeleton 30 (8.1%).

**Figure 12:**
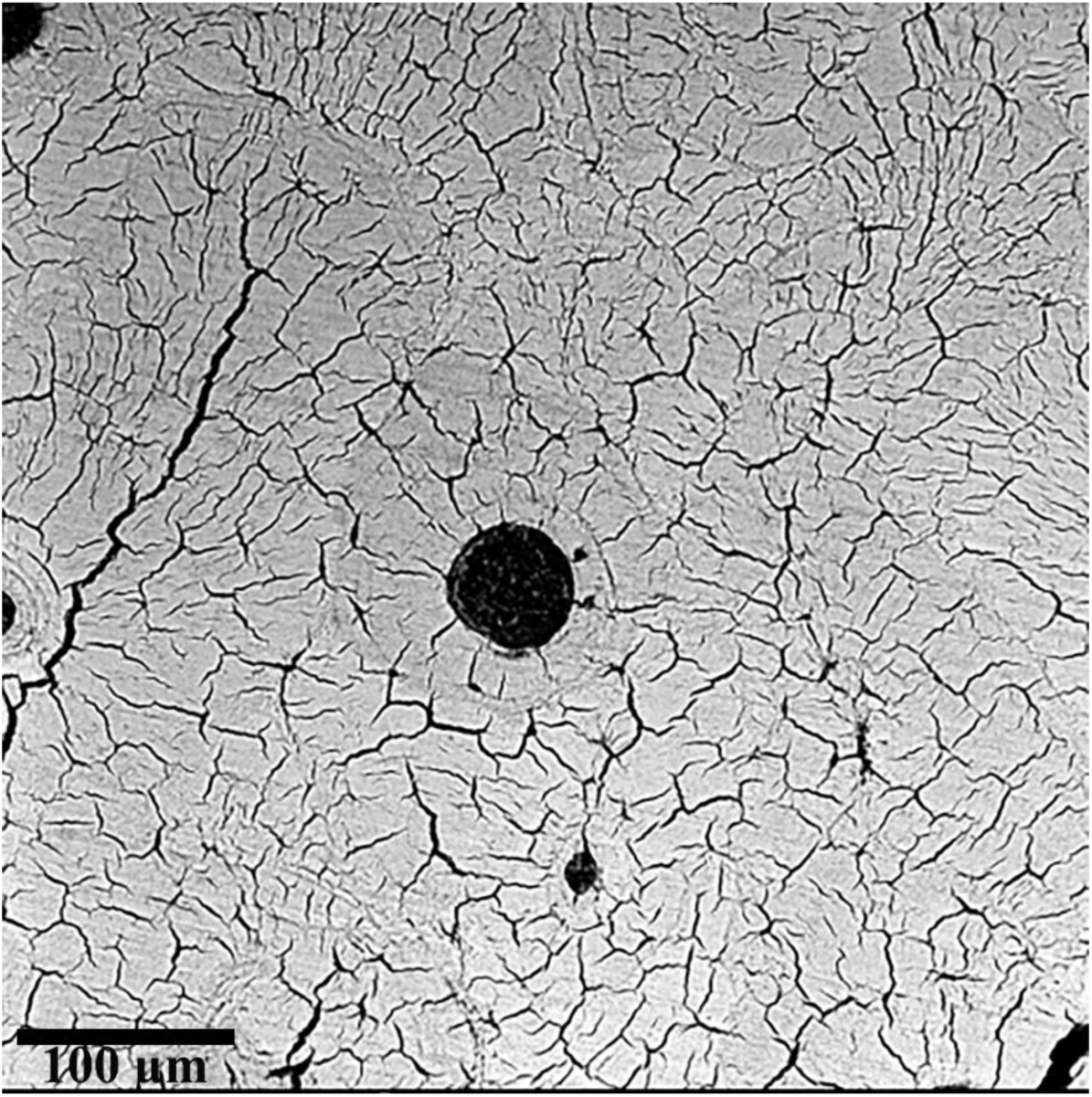
Microcracks observed with SEM on the rib from Object 34_skeleton 1.

### 3.6. Correlations

#### 3.6.1. Percentage of bone preservation

A positive correlation was found between burial depth and the percentage of bone preservation, with deeper burials displaying better levels of preservation (Spearman’s correlation: rho=0.36, S=26762, p=0.004). This pattern was also reflected in differences between the medieval and Kokel burials; the former generally had shallower burial depths, thus exhibiting lower levels of bone preservation than the latter (Mann-Whitney U test: W=318.5, p=0.01). However, this relationship was likely confounded by the uneven distribution of bone type and the use of coffins across burial depths. As shown in Figure 13a, samples from the deepest burials consisted exclusively of long bones (humerus and femur), while ribs dominated the samples from shallower contexts. A similar pattern was evident for coffin use: non-coffin burials occur only in the upper half of the burial depth range, while deeper burials are consistently associated with coffins (Figure 13b). Although both medieval and Kokel contexts included shallow non-coffin burials, all medieval burials lacked coffins. Additional potential confounding factors included the concentration of immature individuals in rib samples (22 of 24 immature individuals are represented by rib samples) and a strong association between coffin use and iron objects: 46 of 48 coffin burials contain iron objects (96%), whereas 22 of 25 non-coffin burials lacked iron (88%). As a result, it was difficult to disentangle the relative contributions of burial depth, age group, bone type, coffin use and associated material culture, and the observed correlation in bone preservation could not be attributed to any singular variable.

**Figure 13:**
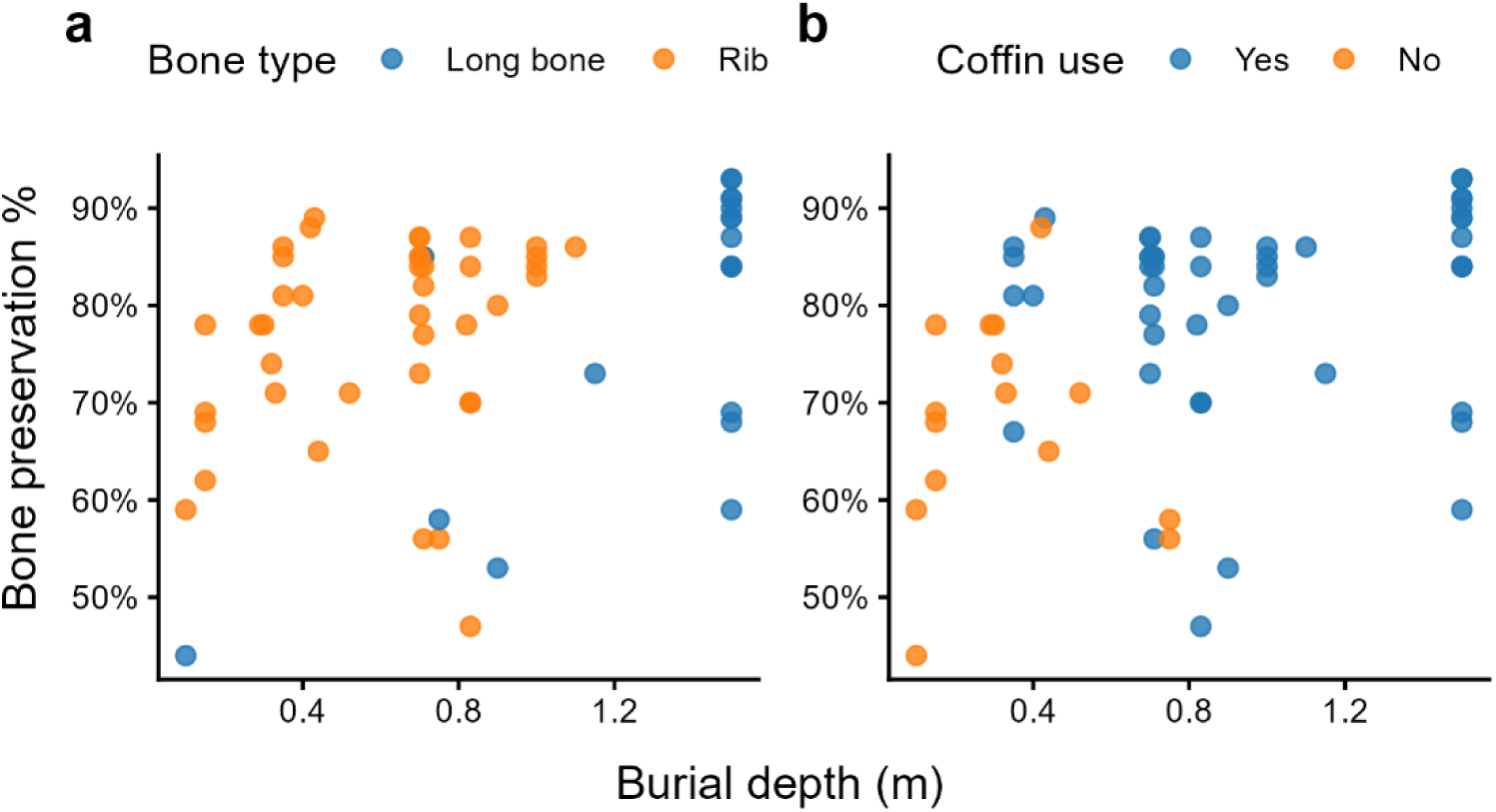
Distribution of quantitative bone preservation percentage by burial depth in relation to sample bone type (a) and coffin use (b).

Looking at the age-at-death of the individuals, no significant difference in bone preservation was observed among the three broad age groups (Kruskal-Wallis test: X^2^=3.91, df=2, p=0.14). Similarly, no significant difference was noticed between sexes of the adult burials in cases where sex could be determined (n=32) (Mann-Whitney U test: U=95, p=0.55).

#### 3.6.2. Percentage of overall cracking

Looking at Figure 14a, samples with higher bone preservation more consistently exhibit lower cracking percentages, while those with lower bone preservation show more variation in overall cracking abundance. However, a Spearman’s correlation test indicates no significant correlation between these two metrics (rho=-0.20, S=80908, p=0.09). Also, in contrast to bone preservation percentage, overall cracking shows no statistical relationship between burial depth (Spearman’s correlations: rho=-0.05, S=43699, p=0.70) or coffin use (Mann-Whitney U test: U=483.5, p=0.18).

**Figure 14:**
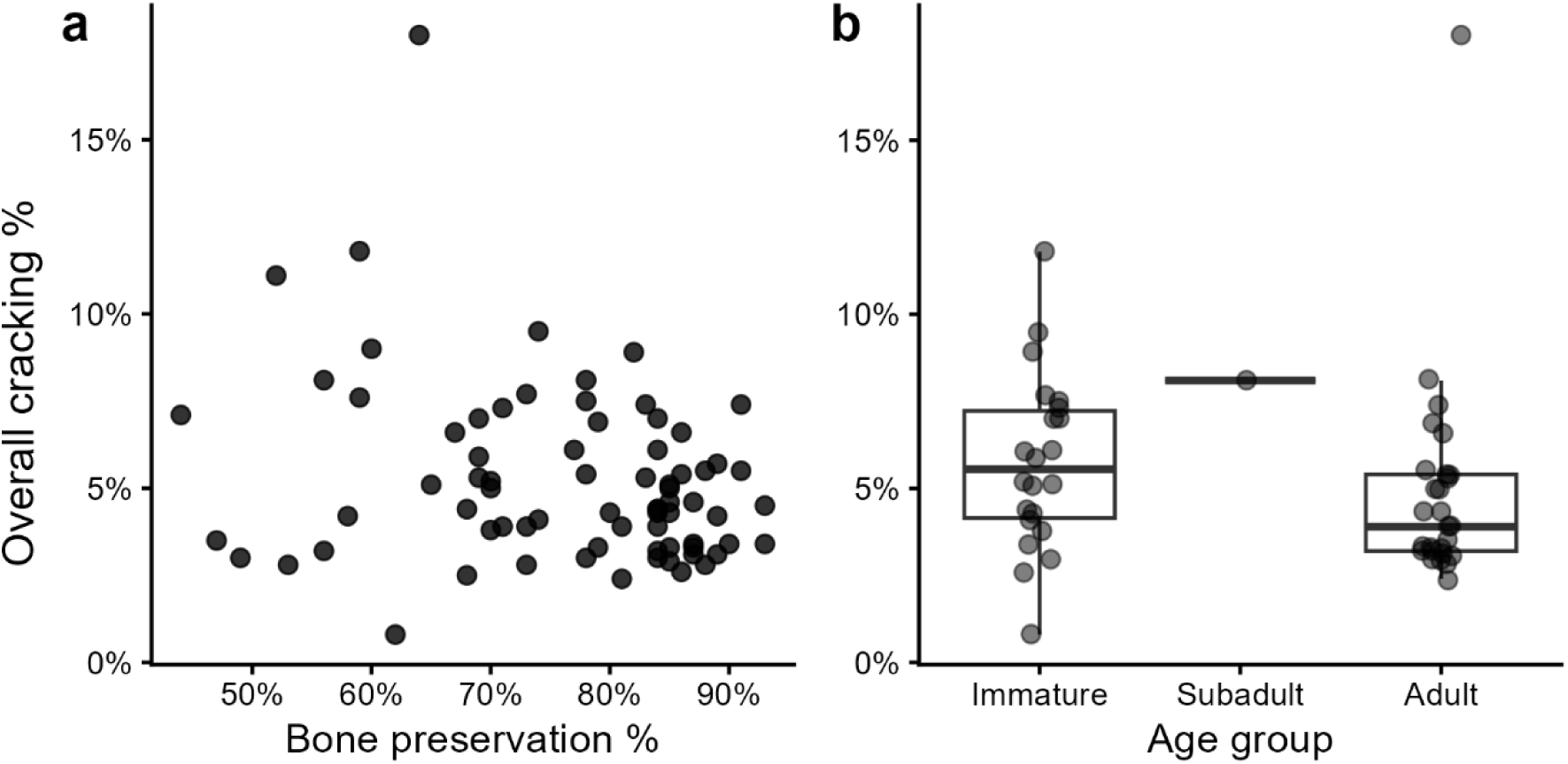
Distribution of overall cracking percentage by bone preservation for all samples (a), and by age group in rib samples only (b).

In terms of biometric variables, most of the samples from the immature age group are represented by rib bones (22 of 24), while the adult samples have a more even distribution of ribs versus long bones (n=27 and 14, respectively). Looking at the adult samples only, there is no clear difference in overall cracking percentage between the two bone types (Mann-Whitney U test: U=207.5, p=0.62). Similarly, no difference is detected between sexes in the adult samples where sex was determined (Mann-Whitney U test: U=124.5, p=0.57). If we consider only the rib samples, the immature group appears to exhibit higher overall cracking abundance over the adult group (Figure 14b), although this difference does not reach the critical threshold for statistical significance (Mann-Whitney U test: 392.5, p=0.06). The subadult category was excluded from comparison due to having a sample size of one.

## 4. Discussion

### 4.1. Bone preservation in a frozen environment

Permafrost is known to provide favourable preservation conditions to organic material as microbial activity is slowed or halted due to frozen soils, low temperatures and anoxic conditions (Hollesen et al. 2017; Loktu and Brødholt 2026). For this reason, exceptionally well-preserved remains, such as frozen mummies, are often recovered from permafrost contexts (Boeskorov et al. 2013; Papageorgopoulou et al. 2015; Slepchenko et al. 2019; Piombino-Mascali and Carr 2021; Slepchenko et al. 2021). Calábková et al. (2023) illustrated this preservation effect experimentally: a Bison metacarpus kept frozen within permafrost in North Russia until recovery displayed very good preservation, whereas an Equus metacarpus that had remained outside the permafrost for several years, exposed at the surface, exhibited Microscopical Focal Destruction (MFD). The authors linked this contrast to ongoing climate change, whereby skeletal remains are increasingly emerging from cryogenic soils and becoming exposed at the surface, accelerating their degradation. The site of Tunnug 1 illustrates both sides of this preservation effect. Wooden structures at the site remain well-preserved, consistent with the protective influence of the frozen environment (Caspari et al. 2020). However, the histological analysis of the skeletal remains revealed evidence of microbial degradation (OHI 2–4) and bone microcracking. This pattern parallels the surface-exposed Equus specimen described above: the Tunnug 1 burials are no longer situated within continuous permafrost and are instead subject to weathering processes, which is reflected in the presence of MFD in their bone microstructure.

### 4.2. Bone degradation: diagenetic features

Overall, the assemblage from Tunnug 1 displayed moderate (OHI 2-3) to good (OHI 4) bone preservation, with the majority of samples scoring 3. This level of preservation is broadly consistent with findings from other archaeological sites in cold or waterlogged environments, where anoxic and stable depositional conditions are known to limit diagenetic activity (Hedges 2002; Kendall et al. 2018). Six individuals, however, exhibited more significant bone degradation: Skeleton 5, Skeleton 12, Skeleton 17, Skeleton 30, Object 23 and Object 51-1. These individuals belonged to different age groups and were buried across different periods and locations within the site (S1, S2 and Figure 3) yet shared a number of funerary characteristics: burial within stone settings and without coffins or iron objects.

The occurrence of MFD at Tunnug 1 was restricted to only three individuals, despite the majority of the assemblage sharing broadly similar burial conditions. This limited prevalence may in part reflect the stabilising influence of permafrost, which is known to restrict microbial activity by maintaining consistently low temperatures and limiting the availability of liquid water necessary for bacterial proliferation (Hedges 2002; Kendall et al. 2018). The fact that Skeletons 12 and 17 were buried in close proximity to one another, while Skeleton 5 was located further south within the funerary area (Figure 3), may point to localised variation in soil conditions, moisture levels or permafrost depth as contributing factors. All three individuals were likely from the Kokel chronology and shared similar funerary treatment, single pits filled with stones, without coffins or grave goods, yet other individuals buried under comparable conditions showed no evidence of MFD. The precise reasons for this disparity remain unclear and most likely reflect the multifactorial and site-specific nature of biological diagenesis (Hollund et al. 2012; Turner-Walker et al. 2023).

Skeleton 30 presented distinct round holes confined to the pleural surface of the rib, suggesting localised bone loss of possible chemical origin. The restricted distribution of these features, concentrated in a single area rather than distributed throughout the section, may reflect localised variation in the depositional microenvironment rather than a systemic diagenetic process. Hedges (2002) noted that increased porosity and dissolution are typically associated with environments subject to water activity, while Turner-Walker and Jans (2008) described how fluctuations in local hydrology can produce non-Wedl diagenetic features through dissolution and demineralisation. Both processes are plausible at Tunnug 1, given the documented proximity to the river, the waterlogged soil conditions and the influence of seasonal freeze-thaw cycles on groundwater movement.

It should be acknowledged, however, that definitively classifying these features as chemical rather than biological degradation is difficult, as similar rounded voids can result from both processes. Micro-CT and SEM analyses did not provide sufficient resolution to resolve this ambiguity conclusively.

Enlarged canaliculi were observed in every sample from Tunnug 1, making them the most consistently present diagenetic feature across the entire assemblage. However, their interpretation remains problematic due to ongoing terminological and definitional inconsistencies in the literature. Some researchers classify enlarged canaliculi as Wedl tunnelling type 2, implying a biological origin (Trueman and Martill 2002; Jans 2005; Brönnimann et al. 2018; Haddow et al. 2023), while others use the term descriptively without specifying how enlargement was measured or what threshold distinguishes enlarged from normal canaliculi (White and Booth 2014; Mavroudas et al. 2023; Mein and Williams 2023). This lack of standardisation makes it difficult to compare observations across studies or to draw meaningful interpretive conclusions from their presence alone.

In the context of Tunnug 1 specifically, it is worth noting that enlargement of canaliculi has been attributed to the mechanical pressure exerted by ice crystal formation during freezing, as expanding water within the canalicular network may widen these spaces over repeated freeze– thaw cycles (Jans 2005; Tersigni 2007). Given the permafrost conditions and long-term climatic fluctuations documented at the site, this mechanism cannot be excluded as a contributing factor. The presence of enlarged canaliculi in all samples, including those with otherwise good bone preservation, is consistent with this interpretation, but need to be confirmed by more targeted analysis. This ambiguity further underscores the need for standardised definitions and measurement criteria for enlarged canaliculi in future histotaphonomic research.

SEM-EDX analysis identified two sources of staining in the bone samples: soil infiltration, evidenced by the presence of silicon and aluminium and iron infiltration originating from both the burial soil and, in some cases, iron grave goods. The external surfaces of the bones, periosteal and endosteal for cortical elements and pleural and cutaneous for the ribs, were most consistently affected, reflecting the inward progression of infiltration from the burial environment.

The observation that iron-stained areas appeared less degraded than the surrounding bone matrix is noteworthy (Figure 5). This pattern was particularly evident in Skeleton 5, where the periosteal surface, heavily infiltrated by iron, was markedly better preserved than the remainder of the femoral cortex. This is consistent with the proposed antibacterial properties of iron, whereby metallic ions released during corrosion may inhibit microbial activity in the immediately surrounding microenvironment (Janaway 1996; Owsley and Compton 1997). Although no statistically significant relationship was identified between the presence of iron grave good items and overall bone preservation scores, the localised protective effect observed at the bone surface level suggests that iron infiltration, whether from artefacts or from the soil itself, may nonetheless exert a taphonomic influence that aggregate scoring methods are insufficiently sensitive to detect.

Object 23 and Object 51-1 present both similarities (no coffin and no grave goods) and major differences: they were buried in different locations within the site and belong to different periods and age groups (Object 23: an adult from the Kokel period; Object 51-1: a neonate from the medieval period). Despite these differences, both presented a combination of rounded holes in the bone structure and hyper-and demineralised areas potentially consistent with MFD, suggesting that multiple diagenetic processes may have operated simultaneously or sequentially within the same individual. Multiple degradation types co-occurring in the same samples have already been documented elsewhere (Papakonstantinou et al. 2020; Loy et al. 2023).

This co-occurrence of features separately attributable to chemical dissolution and bacterial bioerosion complicates classification and interpretation, as it is not always possible to determine whether these represent distinct, independent processes or whether one facilitated the other. For instance, through prior chemical weakening of the bone matrix, creating pathways for bacterial infiltration. Notably, other individuals at the site share similar burial characteristics (absence of coffin and grave goods) without displaying comparable diagenetic features, which further underscores the difficulty of predicting specific diagenetic outcomes from funerary variables such as time period, site location, or age class. Taken together, these observations reinforce the widely acknowledged multifactorial nature of bone diagenesis (Hedges 2002; Kendall et al. 2018; Turner-Walker et al. 2023) and the difficulty of assigning singular causal explanations to diagenetic features in complex archaeological contexts.

Part of this complexity may, however, be partly explained by individual factors: Object 51-1 was a neonate, whose inherently high bone porosity and lower mineralisation may have rendered the bone more susceptible to multiple forms of degradation simultaneously. The particular vulnerability of immature skeletal material to diagenetic alteration is discussed below.

### 4.3. Bone cracking

Cracking percentages ranged from 1% to 18% across the assemblage, with the most heavily cracked samples corresponding to those exhibiting the greatest degree of bone degradation, notably Skeleton 5 (18% and 11.1%), Object 51-1 (11.8%) and Skeleton 30 (8.1%). This pattern suggests that biological and chemical diagenesis weakens the bone matrix, rendering it more susceptible to crack formation, a relationship that has been noted in other histotaphonomic studies (Kendall et al. 2018).

Previous studies about bone cracking in frozen environments (Pokines et al. 2016; Turpin 2017; Trenchat et al. under review) identified ‘long black cracks’ as a result of the freezing cycles on bone microstructure. Those features were not observed for the Tunnug 1 samples. Several factors may account for this absence. First, buried remains are subject to soil pressure and constrained microenvironments that may prevent the unrestricted expansion associated with freeze–thaw cycling in surface-deposited material. Second, the stabilising influence of permafrost, which maintains consistently low temperatures and limits repeated cycles of freezing and thawing, may reduce the mechanical stress associated with ice crystal formation within the bone. Third, centuries of diagenesis, soil compaction and post-depositional disturbance may have obscured or altered crack morphologies that were originally present. The absence of long cracks at Tunnug 1, therefore, does not necessarily preclude a freeze–thaw influence on the bone microstructure but rather highlights the fundamental difference between experimental modern contexts and complex long-term archaeological depositional environments.

Microcracks located on and around the osteonal structures were, however, present in all samples and were visible exclusively under SEM. Similar features have been documented in modern surface-deposited human remains (Schotsmans et al. 2024), experimentally cold-exposed faunal bone (Turpin 2017) and fossilised Jurassic bone (Pfretzschner and Tütken 2011). Their consistent presence across such diverse contexts suggests that they may reflect a combination of *in vivo* biomechanical loading, diagenetic processes and possibly freeze–thaw related microstructural stress, though their precise origin at Tunnug 1 cannot be determined from the current data alone.

### 4.4. Differences between samples and individuals

#### 4.4.1. Intra-individual variations

The marked difference in bone preservation between the rib and femur of Object 23, with the rib exhibiting substantially greater degradation, confirms that skeletal elements from the same individual can undergo divergent taphonomic histories. This reflects differences in intrinsic bone properties, such as cortical thickness, density or trabecular structure, which can influence susceptibility to diagenetic alteration (Hedges 2002; Kendall et al. 2018; Schotsmans et al. 2024). Localised variation in the depositional microenvironment around different parts of the body, such as differential soil moisture, pH or proximity to organic material, may also have contributed. Unfortunately, no field observations documenting differences in soil conditions or state of preservation between the upper and lower body of Object 23 were recorded during excavation, limiting the interpretive possibilities. This highlights the importance of systematic taphonomic recording during fieldwork, as contextual information gathered at the time of excavation is often irreplaceable for post-excavation analyses.

#### 4.4.2. Bone degradation in immature individuals

The microstructural characteristics of immature bone, including lower mineralisation, higher porosity, and the predominance of primary lamellar rather than secondary remodelled bone, are well documented and are known to influence diagenetic susceptibility (White and Folkens 2005; Trenchat et al. 2025). These features were evident in the immature samples from Tunnug 1, particularly the pronounced porosity observed in the neonate individuals.

None of the three neonates from Tunnug 1 exhibited tunnels of degradation. However, Object 51-1 showed significant demineralisation and bone loss, indicating that other forms of degradation can affect neonatal bone even in the absence of bacterial bioerosion. The three neonates shared broadly similar burial characteristics: shallow single pits, without coffins or grave goods, yet their preservation varied noticeably, with Object 51-1 being the most severely degraded. This variability, despite comparable burial conditions, is consistent with the findings of Caruso et al. (2021), who demonstrated that immature bone degradation is influenced by a complex interaction of biological and environmental factors that cannot be reduced to burial type alone. The small sample size of three individuals, however, precludes any broader generalisations, and further research on larger assemblages of immature individuals from comparable depositional contexts is needed.

The relationship between immature bone and diagenetic alteration has more broadly been the subject of considerable debate. Studies have reported an absence of bacterial bioerosion in neonates and young infants, attributed to the underdevelopment of the gut microbiome at or shortly after birth (White and Booth 2014; Booth et al. 2016). On this basis, some researchers have proposed that the absence of MFD may serve as a biomarker for distinguishing stillborn individuals or infanticide victims in archaeological assemblages (Booth et al. 2016). However, this interpretation rests on the assumption that MFD originates exclusively from endogenous gut bacteria, a hypothesis that remains highly contested, as discussed in the introduction. The Tunnug 1 neonates do not straightforwardly support or refute this stillbirth/infanticide biomarker model: the absence of MFD across all three is consistent with it, but the marked variation in non-MFD degradation (particularly in Object 51-1) underscores that MFD absence alone cannot be treated as a reliable standalone indicator, given how many other diagenetic pathways can independently affect neonatal bone.

### 4.5. Funerary treatments

Histological and microtomographic analysis revealed no discernible differences in bone microstructure between individuals from single and multiple burials, suggesting that the number of individuals interred within a structure did not systematically influence diagenetic outcomes.

Object 21 presented an unusual funerary arrangement: the individual was placed in a supine flex position, despite the coffin offering sufficient space to accommodate the body in a fully extended supine position. This uncommon positioning choice was made despite there being no spatial constraint requiring it, raising the question of why the body was deliberately flexed within a coffin large enough for an extended placement. This manipulation may suggest either a specific funerary intention or a practical necessity, for example, flexion to ease carrying over long distances, rather than the body being placed directly in its final position immediately following death. The flexion itself is challenging but not inherently difficult to achieve, especially shortly after death or after *rigor mortis* (Schotsmans et al. 2022). However, no distinctive diagenetic features attributable to a delayed or atypical burial process were identified. While it might be hypothesised that a delay between death and burial could alter early post-mortem decomposition dynamics and thus influence subsequent bone diagenesis, no such signal was detectable in the microstructural data.

A second observation is that the looted mass burial of Object 17 did not exhibit diagenetic alterations clearly attributable to anthropogenic disturbance, beyond a slightly elevated degree of iron infiltration. This observation suggests that post-depositional disturbance of this nature does not leave a consistent or readily identifiable microstructural signature, at least within the timescales and conditions represented at Tunnug 1. Collectively, these findings reinforce the well-established principle that funerary treatment and post-depositional history do not translate straightforwardly into specific diagenetic features, underscoring the need for caution when attempting to reconstruct mortuary practices solely based on bone diagenesis (Hollund et al. 2012; Kendall et al. 2018; Turner-Walker 2019, 2023; Schotsmans et al. 2024).

Statistical analysis identified burial depth as a significant predictor of bone preservation, with deeper burials consistently showing higher OHI scores and lower cracking percentages. This pattern is likely attributable to at least two interrelated factors. First, the soil at Tunnug 1 is characterised by waterlogged and anoxic conditions resulting from the combined influence of permafrost and proximity to the river, known to inhibit microbial activity and reduce biological degradation (Hedges 2002; Turner-Walker and Jans 2008; Kendall et al. 2018). Deeper burials would have been more consistently immersed within the permafrost layer itself, where stable low temperatures further limit diagenetic activity. The interaction between burial depth, permafrost depth and groundwater levels at Tunnug 1 is therefore likely to be a key determinant of preservation variability across the assemblage.

The use of wooden coffins, present in the majority of burials, may have provided an additional protective microenvironment by creating enclosed, anoxic conditions that slowed the ingress of soil microorganisms and moisture (Janaway 1996; Owsley and Compton 1997). While coffin use and burial depth are correlated variables that are difficult to disentangle statistically, given the sample size, both are plausible contributors to the preservation patterns observed.

### 4.6. Methodology

This study employed three complementary imaging modalities with different resolutions: micro-CT, transmitted light microscopy and SEM. It is important to differentiate the spatial resolution of an analytic technique, which depends on the type of probes used (e.g. X-ray, transmitted light or electron), from the numerical resolution of the images produced, which determines the level of detail that can be represented and observed within the image. Each method contributed distinct information to the assessment of bone diagenesis due to the different resolutions. Micro-CT (resolution ≥ 10 μm) provided a non-destructive overview of bone preservation across the entire sample volume, enabling the identification of large-scale structural features, such as porosity, soil infiltration and areas of dense mineralisation prior to any invasive preparation. Conducting micro-CT scans before histological sectioning allowed features observed in thin sections to be cross-referenced with the three-dimensional structure of the intact sample, providing a means of distinguishing genuine taphonomic features from artefacts introduced during cutting and polishing. Transmitted light microscopy enabled the assessment of bone microstructure on a higher resolution (≥ 1 μm), the identification of MFD and enlarged canaliculi, and the scoring of the OHI across the full section area. SEM proved important for the detection of microcracks and the characterisation of mineralisation patterns, as its superior resolution (< 1 μm) revealed features that were invisible under light microscopy. These observations are consistent with findings from other studies that have highlighted the limitations of light microscopy alone for microcrack assessment (Tersigni 2007; Schotsmans et al. 2024). The choice of method, therefore, depends on the research question and available resources, but also on the scale of the features of interest (Trenchat et al. 2024).

The consistent direction of discrepancy between Traditional and Quantitative OHI scores, with the Traditional OHI being systematically higher, warrants careful consideration. The Traditional OHI is inherently subjective, relying on the observer’s assessment of the proportion of preserved bone across a section, and is known to be sensitive to section quality and inter-observer variability (Trenchat et al. 2025). The quantitative approach developed by Trenchat et al. (2025) offers improved reproducibility and objectivity, but its accuracy remains contingent on consistent section preparation, a challenge that is particularly acute with fragile or heavily degraded archaeological materials. Neither method alone is therefore sufficient and the combination of both provides a more robust and internally consistent assessment of bone preservation than either could achieve independently. Future refinements to the quantitative method, such as incorporating correction factors for section thickness variations, could further improve its accuracy and applicability to poorly preserved archaeological assemblages.

This study employed three complementary imaging modalities: micro-CT, transmitted light microscopy and SEM. Each method contributed distinct and non-redundant information to the assessment of bone diagenesis. Micro-CT provided a non-destructive overview of bone preservation across the entire sample volume, enabling the identification of large-scale structural features such as porosity, soil infiltration and areas of dense mineralisation prior to any invasive preparation. Crucially, conducting micro-CT scans before histological sectioning allowed features observed in thin sections to be cross-referenced with the three-dimensional structure of the intact sample, providing a means of distinguishing genuine taphonomic features from artefacts introduced during cutting and polishing.

### 4.7. Limitations

The diversity of the Tunnug 1 assemblage, spanning multiple cultural periods, age groups, funerary treatments and depositional contexts, offered considerable analytical potential but also introduced substantial interpretive complexity. While certain diagenetic features could be linked to specific variables, such as burial depth or the absence of coffins, the majority of preservation patterns could not be readily explained by a single taphonomic factor. This reflects the multifactorial nature of taphonomy and bone diagenesis, in which numerous environmental, biological and contextual variables interact in ways that are difficult to isolate or predict (Kendall et al. 2018; Turner-Walker et al. 2023). In addition, even with the diversity of Tunnug 1 samples, some factors investigated are limited by small sample sizes (i.e. three immature individuals, three individuals with MFD, two individuals with holes) and preclude any broader generalisations. Further research on larger assemblages from comparable depositional contexts is needed.

At Tunnug 1 specifically, interpretation challenges are compounded by centuries of climatic fluctuations affecting soil conditions, groundwater levels, permafrost dynamics and river activity, all of which would have varied over the depositional history of the site in ways that cannot be reconstructed from the archaeological record alone. This also serves as a reminder that, as a result of global warming and rising temperatures, previously protected remains are increasingly being exposed at the surface due to the glacier thawing and are therefore in danger. Consequently, further research is needed to develop strategies for protecting these remains in the future (Bourgeois et al. 2007; Caspari et al. 2018). Controlled experimental research, examining specific variables such as freeze–thaw cycles, soil moisture and burial depth under standardised conditions, remains essential for disentangling these influences and improving the interpretive framework for sites of this nature.

Terminological inconsistencies in the diagenesis literature, as illustrated by the case of enlarged canaliculi discussed above, further complicate the interpretation of microstructural features and limit comparability across studies. Establishing standardised definitions and measurement criteria for key diagenetic features would substantially improve the reliability and reproducibility of histotaphonomic analyses.

The archaeological literature on the Kokel culture remains limited due to the large number of unpublished studies, the difficulty of accessing existing publications for international researchers and the lack of widely available research materials (Sadykov et al., 2021; Sadykov et al., 2025). Consequently, a meaningful comparison between the results of Tunnug 1 and those from other sites of the same region and period is challenging. Future interest in expanding archaeological and anthropological research on the Kokel culture, particularly in the preservation of their burial mounds, would be highly valuable.

A further limitation of this study concerns the secondary nature of the samples. Because the excavation team and the team responsible for microstructural analysis were different, contextual observations made during fieldwork, such as soil colour and texture, moisture levels, the precise spatial relationships between skeletal elements and the condition of organic materials, had been insufficiently recorded or lost in the transfer of information. Labelling inconsistencies between object and skeleton numbers further complicated the dataset and introduced uncertainty in the attribution of some samples. These issues highlight the importance of close collaboration between excavation and laboratory teams from the earliest stages of a project and of systematic taphonomic recording protocols during fieldwork that anticipate the needs of subsequent microstructural analyses. The relatively small sample sizes for certain subgroups, particularly medieval individuals and neonates, also limit the conclusions that can be drawn for these categories, and larger targeted sampling strategies would be beneficial in future work at this and comparable sites.

## 5. Conclusion

This histotaphonomic study of the Tunnug 1 assemblage revealed variable but generally moderate bone preservation (OHI 2-4). Six individuals exhibited more significant diagenetic alteration. Despite the diversity of funerary treatments represented at the site, no clear and consistent microstructural signatures could be attributed to specific burial practices. Burial depth emerged as the most significant predictor of preservation, likely reflecting the combined protective influence of anoxic waterlogged soil conditions and proximity to the permafrost layer. However, this relationship was confounded by correlated variables, including bone type, coffin use and the presence of iron grave goods, underscoring the difficulty of isolating single causal factors in archaeological diagenesis. The absence of long freeze–thaw cracks, documented in experimental modern contexts, underscores the fundamental difference between controlled short-term studies and the complex multi-century depositional history of an archaeological site.

With respect to the study’s objectives, differences in microstructural preservation between burial types could not be demonstrated, and the influence of the freezing environment on bone microstructure could not be isolated from the many other interacting taphonomic variables. These findings reflect the inherently multifactorial nature of bone diagenesis and reinforce the need for targeted experimental research under controlled conditions to better characterise the specific effects of freeze–thaw cycles on buried human remains over long timescales.

Several unusual funerary observations, including the atypical flex positioning of Object 21 and the absence of a distinct diagenetic signature associated with the looted burial of Object 17, illustrate that funerary treatment and post-depositional history do not translate predictably into specific microstructural features. This reinforces the need for caution when using bone diagenesis alone to reconstruct mortuary practices, and points instead to the value of integrating histotaphonomic data with broader contextual and archaeological evidence.

From a methodological perspective, the combination of micro-CT, transmitted light microscopy and SEM proved essential, with each method contributing non-redundant information at different scales of resolution. SEM in particular was indispensable for microcrack detection. The systematic discrepancy between Traditional and Quantitative OHI scores highlights the limitations of subjective visual assessment and points to the value of standardised quantitative approaches, provided that section quality can be adequately controlled. In the future, it would be interesting to combine these methods with higher-resolution imaging, such as synchrotron radiation micro-CT, to better understand the structure and organisation of bone degradation.

Ultimately, Tunnug 1 demonstrates both the promise and the complexity of histotaphonomic research in frozen archaeological contexts and highlights the urgent need for experimental and interdisciplinary approaches to fully disentangle the taphonomic variables that shape bone preservation over centuries of climatic and environmental change. As climate change continues to expose previously protected remains at the surface, this study also reinforces the urgency of developing strategies to document and preserve such material before further degradation occurs.

## Author contributions

**Lolita Trenchat:** Conceptualization, Methodology, Formal Analysis, Software, Validation, Investigation, Writing-Original draft. **Gino Caspari**: Resources, Writing-Review and editing. **Marco Milella**: Resources, Writing-Review and editing. **Timur Sadykov**: Resources, Writing-Review and editing. **Jegor Blochin**: Resources, Writing-Review and editing. **Sam C. Lin**: Formal analysis, Writing-Review and editing. **Nicolas Vanderesse**: Methodology, Formal Analysis, Writing-Review and editing. **Eline M. J. Schotsmans**: Conceptualization, Methodology, Formal Analysis, Validation, Investigation, Resources, Funding, Writing-Original draft.

## Data availability statement

All scores of thin sections are presented in this manuscript. Full scans of the sections are available upon request from the corresponding author

## Acknowledgements

We are grateful to the entire excavation team of Tunnug 1. We thank PACEA and Placamat for accommodating the micro-CT scans analyses. The histological work benefited greatly from the valuable advice of Jose Abrantes and Justyna Miszkiewicz. Dominique Tanner and Lloyd White also generously shared their expertise on SEM analysis, for which we are thankful. Finally, we extend our sincere thanks to Gordon Turner-Walker for his valuable insight and guidance on diagenetic features.

## Declaration of generative AI and AI-assisted technologies in the manuscript preparation process

During the preparation of this work, the authors used Microsoft Copilot and Claude (Anthropic) to assist with writing corrections. The authors reviewed and edited all AI-assisted content and take full responsibility for the published article.

