## Supplementary material S1 for "Bone diagenesis in Iron Age Siberia: a histological and microtomographic study of Tunnug 1 (Russia, 2^nd^–4^th^ c. CE)"

**S1: Information about the burial from Tunnug 1. *: Multiple burials, NA: Variable unknown**.

| **Burial** | **Period** | **Depth (m)** | **Pit measure (m)** | **Pit filling** | **Coffin** | **Notes** |
| --- | --- | --- | --- | --- | --- | --- |
| Object 17* | Kokel | 1.5 | 9x5 | Stones and clay | Yes | Looting pit in the centre |
| Object 5* | Kokel | 0.3 | 8x3 | Stones and clay | Yes | Possible solifluction |
| Object 12 | Likely Kokel | 1.15 | 1.15x2.2 | Stones and clay | Yes | Possible solifluction |
| Object 16 | Kokel | 0.3 | 0.5x2.2 | Clay | No |  |
| Object 18 | Kokel | 0.71 | 2.5x1.5 | Stones | Yes | Repeated flooding by underground water |
| Object 19 | Kokel | 0.4 | 2.2x0.5 | Stones | Yes |  |
| Object 21 | Kokel | 0.7 | 2.5-2.8 | Stones | Yes | Large frost crack near pit. Individual in supine and flexed position. |
| Object 22 | Kokel | 0.55-0.78 | 1 | Stones | Yes |  |
| Object 23 | Kokel | 0.75 | 3x5 | Stones and clay | No |  |
| Object 24* | Kokel | 0.83-0.52 | 3x2.5 | Stones | Yes | Altered by natural processes |
| Object 31 | Kokel | 1.1 | 7x2.5 | Stones | Yes |  |
| Object 32 | Kokel | 0.35 | >1 | Stones | Yes |  |
| Object 33* | Kokel | 1.1 | 7x2.5 | Stones | Yes |  |
| Object 34* | Kokel | 1 | 2x2.5 | Stones | Yes |  |
| Object 35 | Indetermined | 0.32 | 1.15x0.35 | Clay | No |  |
| Object 37 | Medieval | 0.32 | 1x1.1 | Clay | No |  |
| Object 38 | Medieval | 0.52 | 0.9x1.2 | Clay | No |  |
| Object 40* | Kokel | 0.35 | 1.6x0.8 | Mixed loam | Yes | Wood cover |
| Obj42* | Kokel | 0.7 | 2.5 | Stones and clay | Yes |  |
| Object 45 | Medieval | 0.44 | 2.3x1.9 | Stones | No |  |
| Object 46 | Kokel | 0.42 | 1.1x2.2 | Stones and clay | No |  |
| Object 48 | Kokel | 0.15 | No pit | NA | No |  |
| Object 48-49 | Kokel | 0.15 | No pit | NA | No |  |
| Object 51 | Medieval | 0.1 | No pit | NA | No |  |
| Object 51-1 | Medieval | 0.1 | No pit | NA | No |  |
| Object 52 | Medieval | 0.15 | No pit | NA | No |  |
| Object 59 | Kokel | 0.15-0.25 | 1.4x1 | Stones | No | Highly disturbed |
| Skeleton 3 | Kokel | NA | NA | Stones | No |  |
| Skeleton 4 | Likely Kokel | NA | NA | Stones | No |  |
| Skeleton 5 | Likely Kokel | NA | NA | Stones | No |  |
| Skeleton 6 | Likely Kokel | NA | NA | Stones | No |  |
| Skeleton 12 | Likely Kokel | NA | NA | Stones | No |  |
| Skeleton 17 | Likely Kokel | NA | NA | Stones | No |  |
| Skeleton 19 | Indetermined | NA | NA | NA | No |  |
| Skeleton 22 | Likely Kokel | NA | NA | Stones | No |  |
| Skeleton 30 | Kokel | 0.29 | 0.9x0.3 | Stones | No |  |
