## Supplementary material S2 for "Bone diagenesis in Iron Age Siberia: a histological and microtomographic study of Tunnug 1 (Russia, 2^nd^–4^th^ c. CE)"

**S2: Information about the individuals from Tunnug 1. Obj: object, Sk: skeleton, M: Male, F: Female, NA: Variable unknown, ✓: presence of peri-mortem trauma and grave goods (*presence of iron objects).**

| **Individual** | **Bone fragment** | **Sex** | **Age class** | **Body position** | **Grave goods** | **Trauma** | **Notes** |
| --- | --- | --- | --- | --- | --- | --- | --- |
| Obj17_1 | Left Humerus | F | Adult | Disturbed burial (robbed or destroyed in the past).  400 human fragments found (MNI 20 individuals) | ✓* | ✓ | Disturbed |
| Obj17_2 | Left Humerus | M | Adult |  |  |  |  |
| Obj17_3 | Left Humerus | NA | Adult |  |  |  |  |
| Obj17_6 | Left Humerus | NA | Child |  |  |  |  |
| Obj17_7 | Left Humerus | M | Adult |  |  |  |  |
| Obj17_8 | Left Humerus | NA | Adult |  |  |  |  |
| Obj17_9 | Left Humerus | NA | Adult |  |  |  |  |
| Obj17_10 | Left Humerus | NA | Adult |  |  |  |  |
| Obj17_11 | Left Humerus | NA | Adult |  |  |  |  |
| Obj17_12 | Left Humerus | F | Adult |  |  |  |  |
| Obj17_13 | Left Humerus | NA | Subadult |  |  |  |  |
| Obj17_14 | Left Humerus | NA | Adult |  |  |  |  |
| Obj17_15 | Left Humerus | NA | Subadult |  |  |  |  |
| Obj17_16 | Left Humerus | NA | Subadult |  |  |  |  |
| Obj5_Sk1 | Right Femur | M | Old adult | Supine position, head westward | ✓* | ✓ | Found between stones at the north part of the object |
| Obj5_Sk7 | Right Femur | M | Middle adult | Supine and extended position, head to the west | ✓* | ✓ | In the centre of the object |
|  | Rib |  |  |  |  |  |  |
| Obj12_Sk14 | Long bone | F | Old adult | Extended lateral position, head to the west, legs slightly bent at the knees | ✓* |  |  |
|  | Rib |  |  |  |  |  |  |
| Obj16_Sk18 | Right rib | M | Old adult |  | ✓ | ✓ |  |
| Obj18_Sk1 | Rib | M | Young adult | Lateral right position, head to the north | ✓* |  |  |
| Obj18_Sk2 | Right Femur | NA | Adolescent | Lateral right position, head to the north | ✓* | ✓ | Arrowhead in situ (lower thoracic) |
|  | Rib |  |  |  |  |  |  |
| Obj18_Sk3 | Rib | NA | Child | Lateral right position, head to the north | ✓* | ✓ |  |
| Obj18_Sk4 | Rib | NA | Child | Lateral right position, head to the north | ✓* | ✓ | Arrowhead in situ (lower thoracic) |
| Obj18_Sk5 | Rib | NA | Infant | Lateral right position, head to the north | ✓* |  |  |
| Obj19 | Rib | M | Young adult | Head to the northwest | ✓* | ✓ |  |
| Obj21 | Rib | F | Old adult | Supine hyperflexed position | ✓* |  |  |
| Obj22 | Rib | F | Young adult | Extended supine position, head toward the west | ✓* |  |  |
| Obj23 | Right Femur | NA | Young adult |  |  |  |  |
|  | Rib |  |  |  |  |  |  |
| Obj24_Sk1 | Rib | M | Young adult | Supine extended position, head to the north | ✓* | ✓ |  |
| Obj24_Sk2 | Rib | M | Young adult | Supine extended position, head to the north | ✓* | ✓ |  |
| Obj24_Sk3 | Rib | M | Young adult | Supine extended position, head to the north | ✓* | ✓ |  |
| Obj24_Sk4 | Rib | NA | Child | Supine extended position, head to the north | ✓ |  |  |
| Obj24_Sk5 | Rib | NA | Child | Supine extended position, head to the north | ✓* | ✓ |  |
| Obj24_Sk6 | Rib | M | Young adult | Supine extended position, head to the north | ✓* | ✓ |  |
| Obj31 | Rib | F | Old adult | Supine extended position, head to the west | ✓* |  |  |
| Obj32_Sk1 | Rib | M | Old adult | Supine extended position, head to the north-west | ✓* |  | Newborn associated (disturbed) |
| Obj33_Sk1 | Rib | M | Middle adult | Supine extended position, head to the north-west, left arms folded and placed on the right arm | ✓* | ✓ | Arrowhead in situ (left arm) |
| Obj33_Sk2 | Rib | M | Middle adult | Supine extended position, head to the north-west, left arms folded and placed on the right arm | ✓* | ✓ |  |
| Obj34_Sk1 | Rib | NA | Child | Supine extended position, head to the north-west | ✓* | ✓ |  |
| Obj34_Sk2 | Rib | M | Young adult | Supine extended position, head to the north-west, arms folded on the belly | ✓* | ✓ | Arrowhead in situ (upper thorax, right side) |
| Obj34_Sk3 | Rib | M | Young adult | Supine extended position, head to the north-west, arms folded on the belly | ✓* | ✓ | Arrowhead in situ (underneath upper thorax) |
| Obj34_Sk4 | Rib | M | Middle adult | Supine extended position, head to the north-west | ✓* | ✓ | Arrowhead in situ (underneath upper thorax) |
| Obj35 | Rib | NA | Neonate |  |  |  |  |
| Obj37 | Rib | M | Young adult |  |  |  |  |
| Obj38 | Rib | NA | Infant |  |  |  |  |
| Obj40_fauna | Tibia (caprinae) | NA | Young adult |  |  |  |  |
| Obj40_Sk1 | Rib | NA | Child | Extended lateral position, facing Sk2, head toward north-west, feet slightly bent | ✓* | ✓ |  |
| Obj40_Sk2 | Rib | NA | Child | Extended lateral position, facing Sk1, head toward north-west, feet slightly bent | ✓* |  |  |
| Obj42_Sk1 | Rib | F | Middle adult | Supine extended position, head toward the north | ✓* | ✓ |  |
| Obj42_Sk2 | Rib | NA | Old adult | Supine extended position, head toward the north, left hand on the belly. | ✓* | ✓ | Head missing but a knife and a sheep vertebra instead |
| Obj42_Sk3 | Rib | M | Old adult | Supine extended position, head toward the north, left hand on the belly | ✓* |  |  |
| Obj42_Sk4 | Rib | NA | Infant | Near feet Sk1 | ✓* |  |  |
| Obj42_Sk5 | Rib | NA | Child | Near feet Sk2 | ✓* |  |  |
| Obj42_Sk6 | Rib | NA | Infant | Near chest Sk1 | ✓* |  |  |
| Obj45 | Rib | NA | Child |  |  |  |  |
| Obj46 | Rib | M | Middle adult |  |  |  |  |
| Obj48 | Rib | NA | Neonate |  |  |  |  |
| Obj48-49 | Rib | NA | Child |  |  |  |  |
| Obj51_fauna | Left Femur | NA | NA |  |  |  |  |
| Obj51-1 | Rib | NA | Neonate |  |  |  |  |
| Obj52 | Rib | NA | Infant |  |  |  |  |
| Obj59_Sk26 | Rib | NA | Child | Highly disturbed | ✓* |  |  |
| Sk3 | Rib | M | Middle adult | Supine extended position, head to northwest | ✓* |  |  |
| Sk4 | Femur | NA | Child | Incomplete | ✓ |  |  |
|  | Rib |  |  |  |  |  |  |
| Sk5 | Femur | F | Young adult | Fragments of bones | ✓ |  |  |
|  | Rib |  |  |  |  |  |  |
| Sk6 | Rib | NA | Child | Cluster of bones |  |  |  |
| Sk12 | Long bone | NA | Adolescent | Separate bones | ✓ |  |  |
| Sk17 | Tibia | NA | Adolescent | Separate bones |  |  |  |
| Sk19 | Rib | NA | Infant |  |  |  |  |
| Sk22 | Rib | M | Young adult | Upper part in the stone layer, lower part in the natural soil layer | ✓* |  |  |
| Sk30 | Rib | F | Old adult |  |  |  |  |
