## Supplementary material S3 for "Bone diagenesis in Iron Age Siberia: a histological and microtomographic study of Tunnug 1 (Russia, 2^nd^–4^th^ c. CE)"

**S3:** Traditional OHI, Bone preservation quantification (in %) and Quantitative OHI scores for thin sections, corresponding slice from micro-CT scans and entire volume scanned from micro-CT scans. Thin section thickness in μm.

| Individual | Traditional OHI on thin section | Bone preservation on thin section (%) | Quantitative OHI on thin sections | Thin section thickness (in μm) | Traditional OHI on the slide | Bone preservation on the corresponding slice (%) | Quantitative OHI on the corresponding slice | Traditional OHI on the entire scan | Bone preservation on the entire scan (%) | Quantitative OHI on the entire scan |
| --- | --- | --- | --- | --- | --- | --- | --- | --- | --- | --- |
| Obj 17_1 | 4 | 80 | 3 | 116 | 4 | 90 | 4 | 4 | 93 | 4 |
| Obj 17_2 | 3 | 40 | 2 | 229 | 4 | 82 | 3 | 4 | 93 | 4 |
| Obj 17_3 | 3 | 63 | 3 | 129 | 3 | 50 | 3 | 3 | 68 | 3 |
| Obj 17_6 | 3 | 40 | 2 | 104 | 3 | 91 | 4 | 3 | 84 | 3 |
| Obj 17_7 | 3 | 59 | 3 | 203 | 3 | 55 | 3 | 3 | 59 | 3 |
| Obj 17_8 | 3 | 51 | 3 | 148 | 3 | 92 | 4 | 3 | 84 | 3 |
| Obj 17_9 | 3 | 63 | 3 | 153 | 3 | 77 | 3 | 3 | 87 | 4 |
| Obj 17_10 | 3 | 60 | 3 | 173 | 3 | 91 | 4 | 3 | 91 | 4 |
| Obj 17_11 | 4 | 68 | 3 | 202 | 3 | 73 | 3 | 3 | 69 | 3 |
| Obj 17_12 | 3 | 55 | 3 | 167 | 3 | 82 | 3 | 3 | 84 | 3 |
| Obj 17_13 | 4 | 67 | 3 | 122 | 3 | 89 | 4 | 3 | 89 | 4 |
| Obj 17_14 | 4 | 67 | 3 | 106 | 3 | 89 | 4 | 3 | 91 | 4 |
| Obj 17_15 | 3 | 25 | 2 | 108 | 3 | 88 | 4 | 3 | 89 | 4 |
| Obj 17_16 | 3 | 71 | 3 | 211 | 4 | 90 | 4 | 4 | 90 | 4 |
| Obj 5_sk7 (tun7)_rib | 2 | 46 | 2 | 198 | 2 | 54 | 3 | 2 | 53 | 3 |
| Obj 5_sk7 (tun7)_long bone | 2 | 51 | 3 | 200 | 3 | 86 | 4 | 3 | 80 | 3 |
| Obj 12 sk14 | 3 | 54 | 3 | 118 | 3 | 72 | 3 | 3 | 73 | 3 |
| Obj 16_sk18 | 3 | 57 | 3 | 120 | 3 | 86 | 4 | 3 | 78 | 3 |
| Sk3 | 3 | 64 | 3 | 97 | 3 | 78 | 3 | 3 | 83 | 3 |
| Sk4_rib | 3 | 86 | 4 | 108 | no scan | No scan | no scan | no scan | no scan | no scan |
| Sk4_long bone | 3 | 87 | 4 | 170 | no scan | No scan | no scan | no scan | no scan | no scan |
| Sk5_rib | 2 | 15 | 2 | 177 | 2 | 74 | 3 | 2 | 64 | 3 |
| Sk5_long bone | 2 | 30 | 2 | 174 | 2 | 52 | 2 | 2 | 52 | 3 |
| Sk6 | 3 | 69 | 3 | 112 | no scan | No scan | no scan | no scan | no scan | no scan |
| Sk12 | 3 | 54 | 3 | 107 | 3 | 72 | 3 | 3 | 73 | 3 |
| Sk17 | 2 | 19 | 2 | 132 | 2 | 63 | 3 | 2 | 60 | 3 |
| Sk19 | 3 | 37 | 2 | 145 | 3 | 78 | 3 | 3 | 74 | 3 |
| Sk22 | 3 | 58 | 3 | 136 | 3 | 89 | 4 | 3 | 88 | 4 |
| Obj 59_sk26 | 3 | 50 | 3 | 163 | 3 | 77 | 3 | 3 | 78 | 3 |
| Sk30 | 2 | 50 | 3 | 174 | 2 | 72 | 3 | 2 | 78 | 3 |
| Obj 18_Sk1 | 3 | 52 | 3 | 183 | 3 | 85 | 4 | 3 | 85 | 4 |
| Obj 18_Sk2 | 3 | 63 | 3 | 145 | 3 | 86 | 4 | 3 | 85 | 4 |
| Obj 18 Sk2 | 3 | 48 | 2 | 158 | 3 | 63 | 3 | 3 | 56 | 3 |
| Obj 18_Sk3 | 3 | 72 | 3 | 231 | 3 | 72 | 3 | 3 | 82 | 3 |
| Obj 18_Sk4 | 3 | 55 | 3 | 120 | 3 | 79 | 3 | 3 | 77 | 3 |
| Obj 18_Sk5 | 3 | 53 | 3 | 130 | 3 | 84 | 3 | 3 | 84 | 3 |
| Obj 19 | 3 | 65 | 3 | 206 | 4 | 89 | 4 | 4 | 89 | 4 |
| Obj 21 | 3 | 52 | 2 | 278 | 4 | 85 | 4 | 4 | 87 | 4 |
| Obj 22 | 3 | 58 | 3 | 184 | 3 | 80 | 3 | 3 | 78 | 3 |
| Obj 23_rib | 2 | 25 | 2 | 189 | 2 | 59 | 3 | 2 | 56 | 3 |
| Obj 23_femur | 3 | 55 | 3 | 181 | 3 | 66 | 3 | 3 | 58 | 3 |
| Obj 24_Sk1 | 3 | 50 | 3 | 199 | 3 | 87 | 4 | 3 | 87 | 4 |
| Obj 24_Sk2 | 3 | 71 | 3 | 174 | 3 | 81 | 3 | 3 | 70 | 3 |
| Obj 24_Sk3 | 2 | 39 | 2 | 234 | 3 | 86 | 4 | 3 | 84 | 3 |
| Obj 24_Sk4 | 2 | 39 | 2 | 166 | 3 | 73 | 3 | 3 | 70 | 3 |
| Obj 24_Sk5 | 3 | 58 | 3 | 217 | 3 | 80 | 3 | 3 | 70 | 3 |
| Obj 24_Sk6 | 3 | 48 | 2 | 246 | no scan | no scan | no scan | no scan | no scan | no scan |
| Obj 31 | 3 | 55 | 3 | 182 | 3 | 81 | 3 | 3 | 81 | 3 |
| Obj 32_Sk1 | 3 | 59 | 3 | 174 | 3 | 82 | 3 | 3 | 81 | 3 |
| Obj 33_Sk1 | 3 | 49 | 2 | 170 | 3 | 86 | 4 | 3 | 86 | 4 |
| Obj 33_Sk2 | 3 | 42 | 2 | 182 | 3 | 80 | 3 | 3 | 79 | 3 |
| Obj 34_Sk1 | 3 | 59 | 3 | 183 | 3 | 82 | 3 | 3 | 84 | 3 |
| Obj 34_Sk2 | 3 | 64 | 3 | 103 | 3 | 87 | 4 | 3 | 86 | 4 |
| Obj 34_Sk3 | 3 | 75 | 3 | 191 | 3 | 83 | 3 | 3 | 83 | 3 |
| Obj 34_Sk4 | 3 | 46 | 2 | 314 | 3 | 85 | 4 | 3 | 85 | 4 |
| Obj 35 | 2 | 20 | 2 | 182 | 3 | 76 | 3 | 3 | 74 | 3 |
| Obj 37 | 3 | 56 | 3 | 232 | 3 | 78 | 3 | 3 | 71 | 3 |
| Obj 38 | 2 | 26 | 2 | 199 | 3 | 70 | 3 | 3 | 71 | 3 |
| Obj 40 | 3 | 67 | 3 | 251 | no scan | no scan | no scan | no scan | no scan | no scan |
| Obj 40_Sk1 | 3 | 58 | 3 | 210 | 3 | 86 | 4 | 3 | 85 | 4 |
| Obj 40_Sk2 | 3 | 54 | 3 | 153 | 3 | 82 | 3 | 3 | 86 | 4 |
| Obj 42_Sk1 | 3 | 63 | 3 | 208 | 3 | 85 | 4 | 3 | 85 | 4 |
| Obj 42_Sk2 | 3 | 44 | 2 | 235 | 3 | 85 | 4 | 3 | 85 | 4 |
| Obj 42_Sk3 | 3 | 68 | 3 | 152 | 3 | 80 | 3 | 3 | 79 | 3 |
| Obj 42_Sk4 | 2 | 39 | 2 | 184 | 3 | 75 | 3 | 3 | 73 | 3 |
| Ob 42_Sk5 | 2 | 38 | 2 | 279 | 3 | 89 | 4 | 3 | 87 | 4 |
| Obj 42_Sk6 | 2 | 42 | 2 | 145 | 3 | 84 | 3 | 3 | 84 | 3 |
| Obj 45 | 3 | 60 | 3 | 214 | 3 | 63 | 3 | 3 | 65 | 3 |
| Obj 46 | 3 | 53 | 3 | 193 | 3 | 87 | 4 | 3 | 88 | 4 |
| Obj 48 | 2 | 17 | 2 | 244 | 3 | 73 | 3 | 3 | 68 | 3 |
| Obj 48-49 | 3 | 65 | 3 | 83 | 3 | 64 | 3 | 3 | 62 | 3 |
| Obj 51 | 2 | 44 | 2 | 208 | no scan | no scan | no scan | no scan | no scan | no scan |
| Obj 51-1 | 2 | 29 | 2 | 239 | 2 | 58 | 3 | 2 | 59 | 3 |
| Obj 52 | 2 | 39 | 2 | 204 | 3 | 75 | 3 | 3 | 69 | 3 |
