## Supplementary material S4 for "Bone diagenesis in Iron Age Siberia: a histological and microtomographic study of Tunnug 1 (Russia, 2^nd^–4^th^ c. CE)"

**S4: R Markdown file used for the statistical tests and the Generalised Linear Model (S. Lin)**.

This R Markdown file contains the R code to reproduce the statistical results in “Bone diagenesis in Iron Age Siberia: a histological and microtomographic study of Tunnug 1 (Russia, 2nd–4th c. CE)” by Trenchat and colleagues.

R packages used include:

- ggplot2 (Wickham, H. 2016. ggplot2: Elegant Graphics for Data Analysis. Springer-Verlag, New York.)
- ggpubr (Kassambara A (2026). ggpubr: ‘ggplot2’ Based Publication Ready Plots. doi:10.32614/CRAN.package.ggpubr https://doi.org/10.32614/CRAN.package.ggpubr, R package version 0.6.3, <https://CRAN.R-project.org/package=ggpubr>.)
- ggsci (Xiao N (2025). ggsci: Scientific Journal and Sci-Fi Themed Color Palettes for ‘ggplot2’. doi:10.32614/CRAN.package.ggsci https://doi.org/10.32614/CRAN.package.ggsci, R package version 4.2.0, <https://CRAN.R-project.org/package=ggsci>.)

**Bone preservation percentage**

Spearman’s correlation test between the percentage of bone preservation and burial depth.

```
##
##  Spearman's rank correlation rho
##
## data:  xdata$Pit.depth..in.m. and xdata$Bone.preservation....
## S = 26762, p-value = 0.004004
## alternative hypothesis: true rho is not equal to 0
## sample estimates:
##       rho
## 0.3576627
```

Mann-Whitney U test of bone preservation by chronological period.

```
##
##  Wilcoxon rank sum test with continuity correction
##
## data:  Bone.preservation.... by Chronology_all
## W = 318.5, p-value = 0.01089
## alternative hypothesis: true location shift is not equal to 0
```

Figure 13. Distribution of quantitative bone preservation percentage by burial depth in relation to sample bone type (a) and coffin use (b).

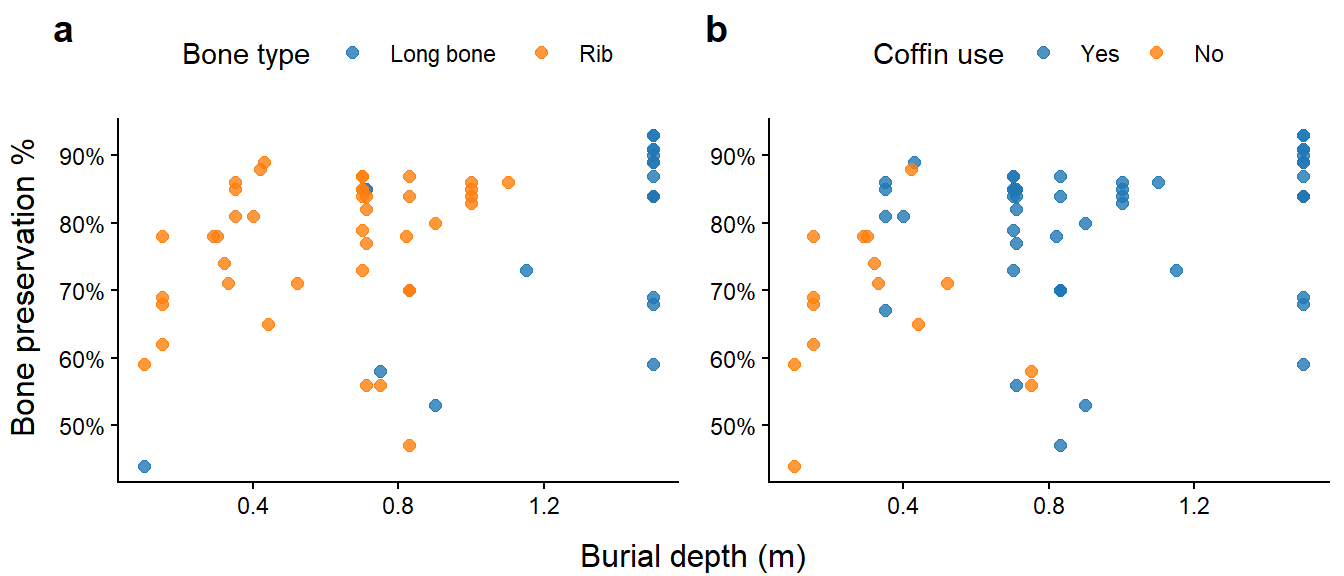

Kruskal-Wallis test of bone preservation by age group.

```
##
##  Kruskal-Wallis rank sum test
##
## data:  Bone.preservation.... by Age_general
## Kruskal-Wallis chi-squared = 3.909, df = 2, p-value = 0.1416
```

Mann-Whitney U test of bone preservation by sex.

```
##
##  Wilcoxon rank sum test with continuity correction
##
## data:  Bone.preservation.... by Sex
## W = 95, p-value = 0.5549
## alternative hypothesis: true location shift is not equal to 0
```

**Overall cracking**

Figure 14. Distribution of overall cracking percentage by bone preservation for all samples (a), and by age group in rib samples only (b).

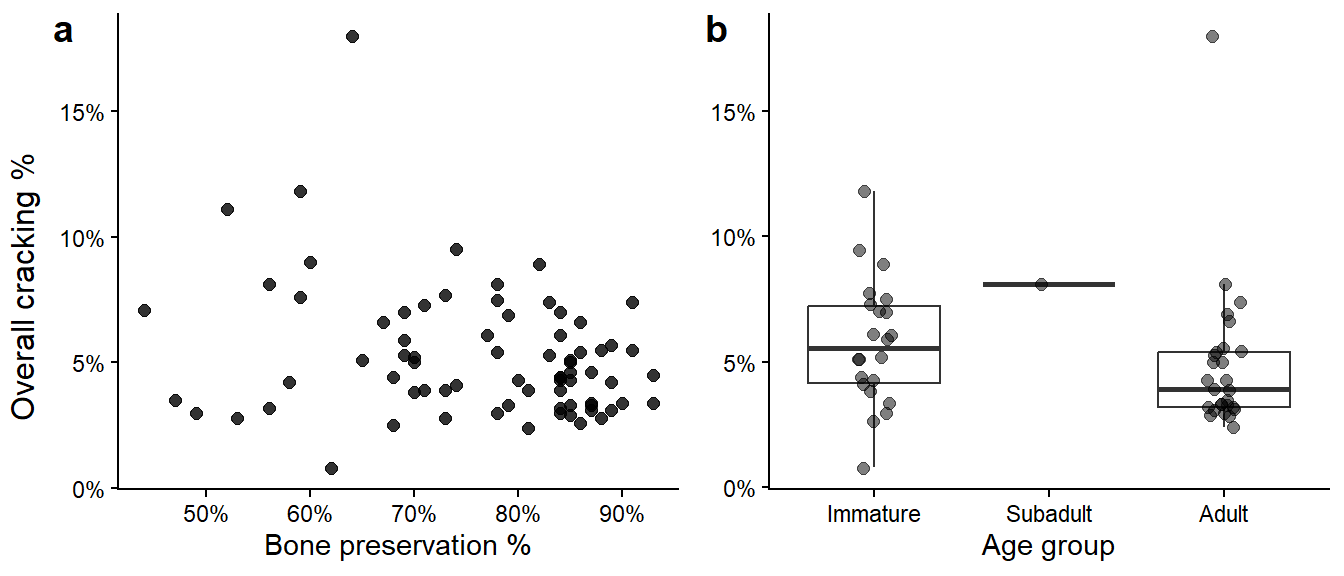

Spearman’s correlation test between the percentage of overall cracking and the percentage of bone preservation.

```
##
##  Spearman's rank correlation rho
##
## data:  xdata$Bone.preservation.... and xdata$Overall.cracking....
## S = 80908, p-value = 0.09052
## alternative hypothesis: true rho is not equal to 0
## sample estimates:
##        rho
## -0.1981875
```

Spearman’s correlation test between the percentage of overall cracking and burial depth

```
##
##  Spearman's rank correlation rho
##
## data:  xdata$Overall.cracking.... and xdata$Pit.depth..in.m.
## S = 43699, p-value = 0.7038
## alternative hypothesis: true rho is not equal to 0
## sample estimates:
##         rho
## -0.04885219
```

Mann-Whitney U test of the percentage of overall cracking by coffin use

```
##
##  Wilcoxon rank sum test with continuity correction
##
## data:  Overall.cracking.... by Coffin
## W = 483.5, p-value = 0.1773
## alternative hypothesis: true location shift is not equal to 0
```

Mann-Whitney U test of the percentage of overall cracking by bone type

```
##
##  Wilcoxon rank sum test with continuity correction
##
## data:  Overall.cracking.... by Bone.type_general
## W = 207.5, p-value = 0.6204
## alternative hypothesis: true location shift is not equal to 0
```

Mann-Whitney U test of the percentage of overall cracking by sex

```
##
##  Wilcoxon rank sum test with continuity correction
##
## data:  Overall.cracking.... by Sex
## W = 124.5, p-value = 0.5687
## alternative hypothesis: true location shift is not equal to 0
```

Mann-Whitney U test of the percentage of overall cracking by age

```
##
##  Wilcoxon rank sum test with continuity correction
##
## data:  Overall.cracking.... by Age_general
## W = 392.5, p-value = 0.05608
## alternative hypothesis: true location shift is not equal to 0
```
